# SmartGate is a spatial metabolomics tool for resolving tissue structures

**DOI:** 10.1101/2022.09.25.509375

**Authors:** Kaixuan Xiao, Yu Wang, Kangning Dong, Shihua Zhang

## Abstract

Imaging mass spectrometry (IMS) is one of the powerful tools in spatial metabolomics for obtaining metabolite data and probing the internal microenvironment of organisms. It has dramatically advanced the understanding of the structure of biological tissues and the drug treatment of diseases. However, the complexity of IMS data hinders the further acquisition of biomarkers and the study of certain specific activities of organisms. To this end, we introduce an artificial intelligence tool SmartGate to enable automatic peak picking and spatial structure identification in an iterative manner. SmartGate selects discriminative m/z features from the previous iteration by differential analysis and employs a graph attention auto-encoder model to perform spatial clustering for tissue segmentation using the selected features. We applied SmartGate to diverse IMS data at multicellular or subcellular spatial resolutions and compared it with four competing methods to demonstrate its effectiveness. SmartGate can significantly improve the accuracy of spatial segmentation and identify biomarker metabolites based on tissue structure-guided differential analysis. For multiple consecutive IMS data, SmartGate can effectively identify structures with spatial heterogeneity by introducing three-dimensional spatial neighbor information.

## Introduction

Spatial metabolomics is a rapidly developing subfield in omics, focusing on deciphering metabolites and some metabolic pathways in spatial context [1-4]. In recent years, the outstanding performance of spatial metabolomics in detecting metabolites and diagnosing diseases has made this field of great interest and dramatically promoted its development [5, 6]. Compared to spatial transcriptomics, metabolites are closer to clinical phenotypes in biological systems and have great potential for identifying molecular mechanisms of disease treatment [7]. Moreover, diverse functional changes such as hormone regulation and intercellular communication of protein macromolecules could be eventually reflected at the metabolic level [7, 8].

Recently, with the usage of imaging mass spectrometry (IMS), spatial metabolomics has been greatly improved [9]. IMS pixelates the surface of the sample, and the mass spectra of each pixel surface were collected. Each pixel has a spectrum where the x-axis represents the mass-to-charge ratio (m/z) and the y-axis represents the relative intensity [10]. New IMS-based techniques like matrix-assisted laser desorption ionization (MALDI), and desorption electrospray ionization (DESI) have been proven promising in many applications like cancer diagnosis and drug development [11]. These technologies profile the relative intensity of metabolites at the multicellular resolution, where the diameter for each pixel is usually between 50μ m-100μm. Recently, some subcellular resolution techniques, such as TOF-SIMS [12] and three-dimensional (3D) OrbiSIMS [13], have been developed. However, the spectrum form and high technical noise of IMS data prevent us from further exploring the possibility of some biological phenomena.

To address the issues of signal noise and high-dimensionality, peak picking is a critical step for preprocessing the IMS data [14], which is usually performed by selecting m/z features with the highest mean values or manually selecting m/z features of interest [15]. However, these methods may ignore the signals of small tissue structures or introduce subjective factors. Thus, how to select discriminative features is one of the great challenges of spatial metabolomics.

Another essential step is identifying tissue structures that are spatially coherent in biomarker metabolites. The most widely used approaches first employ linear dimensionality reduction and then cluster the pixels into groups with the traditional clustering methods, such as K-means and Louvain algorithm. Besides, msiPL [16] adopts a nonlinear deep generative model for dimensionality reduction and a Gaussian mixture model for clustering. However, the above methods are not tailored for spatial metabolomics data and the identified clusters may be spatially discrete due to the lack of consideration of spatial information. To solve this issue, SASA leverages the distance-preserving projection method FastMap to embed the spatial relation between adjacent pixels [17]. SSC introduces the spatially aware distance to regularize the distance between pixel spectrums [18]. SEAM, developed at the subcellular level, first segments nucleus regions leveraging the Potts model and restricted Boltzmann machines by selecting pixels that are highly correlated with nuclear markers. It further represents the metabolic features of each segmented nuclear cell using a distilled softmax space for further clustering [6]. At the same time, several artificial intelligence tools have been developed to identify spatial structures in spatial omics by introducing spatial neighbor graphs to consider spatially correlated structures across tissues [19-21], For example, STAGATE employs a graph attention auto-encoder to integrate the spatial information and gene expression for spatial transcriptomics, showing promising performance in terms of accuracy and computational efficiency [19].

To this end, we aim to simultaneously solve the peak picking and tissue structure identification problems of spatial metabolomics by introducing an iterative graph attention auto-encoder method SmartGate. SmartGate first constructs the spatial neighbor graph based on the relative spatial locations of pixels and enters it into the graph attention auto-encoder with the raw data to obtain the clustering results of the first iteration. Then SmartGate performs automatic peak picking based on the tissue structure-guided differential analysis and uses the selected discriminative features as input to the next iteration to identify more accurate structures. We applied SmartGate to diverse spatial metabolomics datasets profiled by different IMS-based technologies, including Thermo Finnigan LTQ-DESI, MALDI FT-ICR, and nano-DESI IMS and TOF-SIMS, to demonstrate its effectiveness in deciphering tissue structures and identifying biomarker metabolites.

## Results

### Overview of SmartGate

The IMS-based datasets usually consist of about 10^3^∼10^6^ pixels with corresponding spatial coordinates at multicellular or even subcellular resolution for diverse tissue sections [9]. The preprocessed spectrum signals of each pixel are transformed to a metabolic matrix with rows representing pixels and columns representing selected m/z values **(Fig. 1)**. SmartGate first builds a spatial neighbor graph to model the spatial information. SmartGate takes the metabolic matrix and spatial neighbor graph as input and jointly perform the automatic peak picking and tissue segmentation in an iterative manner **(Fig. 1)**. SmartGate first adopts a graph attention autoencoder to extract the original embedding or the representation inspired by STAGATE for the first iteration. SmartGate further uses the encoded representation to identify tissue structures with clustering algorithms, such as mclust [22] and Louvain [23]. At last, SmartGate selects the top m/z features through tissue structure-guided differential analysis as the input for the next iteration until it gets stable features and tissue segmentation.

**Fig. 1.**
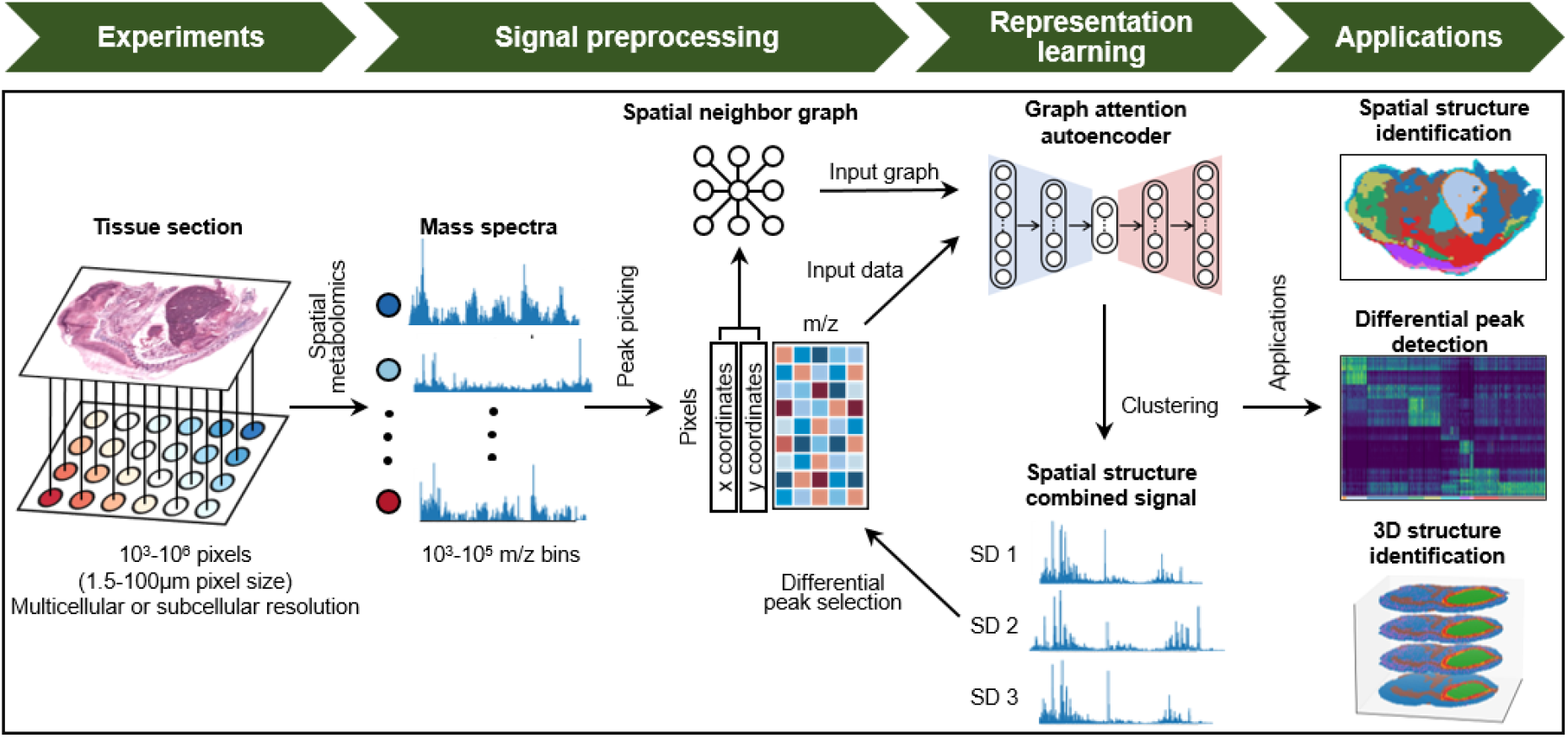
Overview of SmartGate. The IMS data were generated by desorbing the metabolites from tissue sections at the corresponding pixel locations and putting them into the mass spectrometer for analysis, and the original data were subjected to preliminary peak picking to select m/z features of interest. Then, basic peak picking (depending on the data used) is performed to select m/z features, and the spatial information is used to construct a neighbor graph. Both the metabolic information and the spatial neighbor graph are used to reduce the dimensionality of the feature vector by graph attention autoencoder, and the low-dimensional embedding of the feature vector is used for downstream analysis by clustering algorithm for resolving spatial structures. Differential peak analysis is used to select some ions with the most significant differences as the input for the next iteration. The m/z features with the highest discrepancy obtained from the final iteration are used for downstream final analysis, such as spatial segmentation, discrepancy analysis, and spatial segmentation of 3D data.

We applied SmartGate to six spatially resolved metabolomics datasets from diverse platforms and different spatial resolutions and compared it with four methods including GMM, SSC, SASA, and msiPL to demonstrate its effectiveness (**Table 1 and Supplementary Materials**). SmartGate automatically selects potential biologically significant m/z features, providing more accurate information for downstream analysis. SmartGate can significantly improve segmentation accuracy, differential maker identification, and structure identification of three-dimensional (3D) sections.

**Table 1.** Summary of datasets used in this study.

| Dataset | Technology | m/z range (u) | Spatial resolution (μm) | Data size (pixel*peak) | 3D slices or not | Ref |
| --- | --- | --- | --- | --- | --- | --- |
| Pig fetus | Thermo Finnigan LTQ linear ion trap MS -DESI | 150 –1,000 | 100 | 10,200*4,959 | No | [18] |
| PDX mouse brain model of glioblastoma | MALDI FT-ICR IMS | 100-3,000 | 100 | 21,241*3,570 | Yes | [25] |
| Mouse kidney | nano-DESI | 133-2,000 | 50 | 95,000*140 | No | [26] |
| Mouse kidney | MALDI IMS | 2,000-20,000 | 50 | 18,536*7,680 | Yes | [27] |
| Human liver† | TOF-SIMS | 0-2000 | 1.5 | 65,536*228 (*) | No | [6] |
| Mouse liver† | TOF-SIMS | 0-2000 | 1.5 | 65,536*244 (*) | No | [6] |

†The tissue areas of these two slices are 400×400μm<sup>2</sup>
\*indicates the data size after preprocessing (pixel\*peak)

### SmartGate identifies the main structures of the pig fetus dataset

We first applied SmartGate to the pig fetus dataset consisting of 4,959 pixels and 10,200 peaks (selected by experts before) over the range of 150 –1,000 m/z **(Fig. 2a and Table 1)**. SmartGate extracted 3,181 informative m/z features and identified ten spatial structures **(Fig. 2b)**. Compared to GMM, SSC, SASA, and msiPL, SmartGate can not only identify the distinct tissue structures like brain, heart, and liver that are consistent with the ones annotated in the corresponding H&E image **(Fig. 2a)**, but also reveal the sub-structures in the porcine fetal brain, which were segmented as the proencephalon (cluster 8) and mecencephalon (cluster 3) according to a recent study [11] (**Fig. 2c)**. However, GMM, SASA, SSC, and msiPL failed to decipher such substructures for the same number of classes **(Fig. 2b)**. Furthermore, the spatial structure-guide differential analysis found some marker ions that are highly spatially consistent with the identified tissue structures. For example, an ion with an m/z value of 810.53, which was tentatively identified as PS (38:4), was specifically expressed in the proencephalon and mecencephalon tissue substructures **(Fig. 2d, e)**. The high expression of PS lipids that appeared in the brain may be related to apoptosis of the nervous system [11]. Compared to SmartGate, these methods showed more discrete segmentations other than the brain, heart, and liver due to the lack of consideration of spatial information (**Fig. 2c** and **Fig. S1**). Besides, for other regions with significant markers like heart and liver, the result of SmartGate can be verified by several known metabolites, such as the high expression of N-Acetyl-glutamine (m/z=187.08) in heart tissue, which may be related to a certain enzyme in the heart [11], and strong abundance of FA dimer (m/z=535.25) in liver tissue **(Fig. 2d, e)**.

**Fig. 2.**
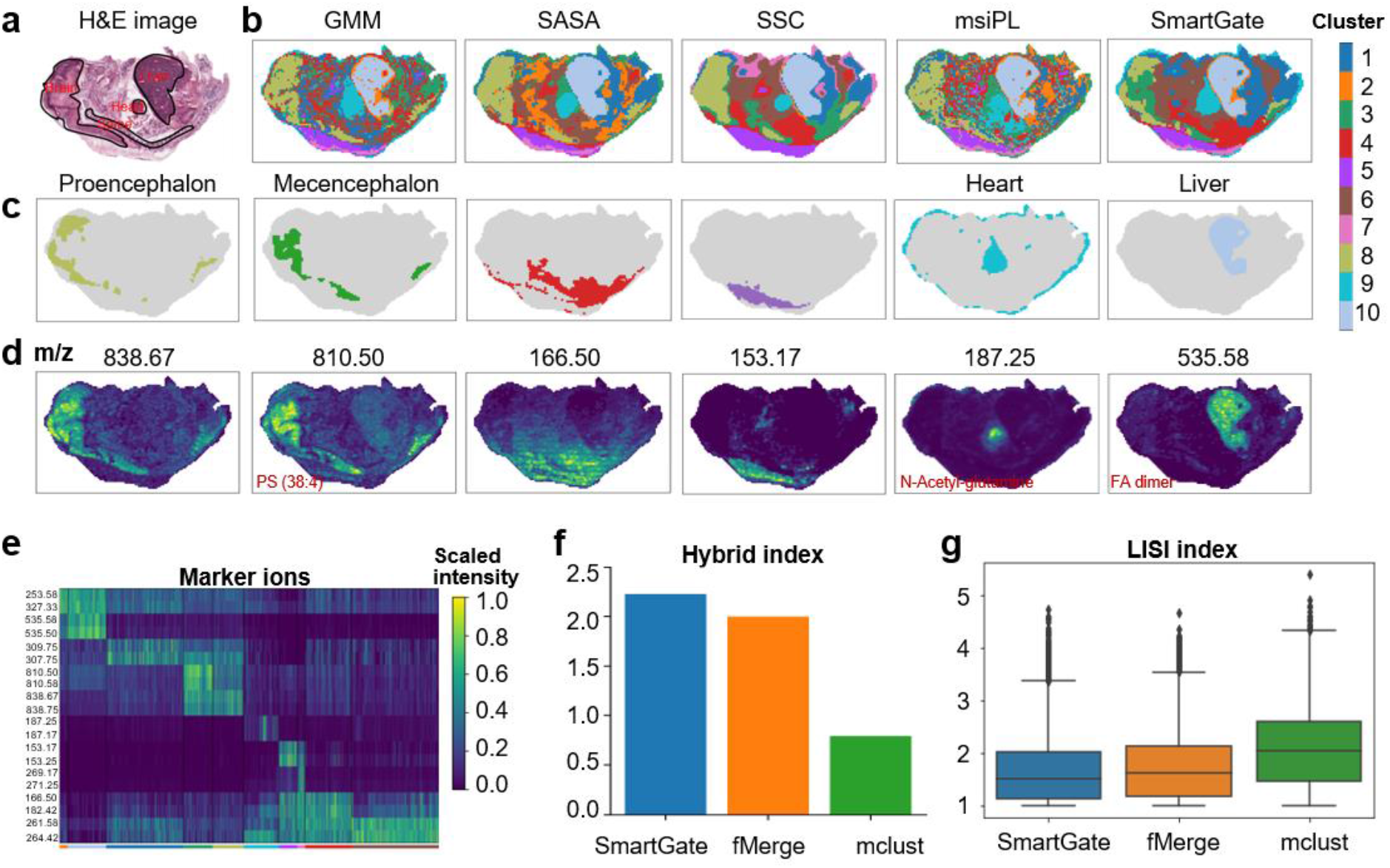
SmartGate identifies the main structures in the pig fetus dataset. **a** The manual annotation of the main structures of the pig fetus in the H&E image. **b** Spatial structures identified GMM, SASA, SSC, msiPL, and SmartGate respectively. **c, d** Spatial distribution and corresponding marker ions of six structures identified by SmartGate. **e** Top marker ions determined by differential analysis of each structure identified by SmartGate. **f, g** Comparison of the Hybrid index score (**f**) and LISI index score (**g**) of SmartGate, fMerge, and mclust respectively.

To illustrate the advantage of SmartGate in automatically selecting m/z features, we compared the clustering results using the features selected by SmartGate, fMerge (mean value taken in sliding window), and mclust-based peak picking process respectively as inputs **(Fig. S2, Materials and Methods)**. The results showed that the tissue structures identified using SmartGate-selected features were more spatially continuous. To verify it, we calculated the hybrid index and local inverse Simpson’s index (LISI) [24] **(Fig. 2f, g, Materials and Methods)** and found that the clustering results using SmartGate-selected features could better capture the potential biological patterns than those using the features selected by fMerge or mclust.

### SmartGate reveals the different metabolic patterns of tumor core and edge

We applied SmartGate to the patient-derived xenograft (PDX) mouse brain model of glioblastoma (GBM) datasets consisting of 3,570 pixels and 21,241 peaks **(Table 1)** [25]. The darker area in the upper side of the H&E image is the transplanted tumor tissue **(Fig. 3a)**. SmartGate and SASA showed better performance in identifying the tumor core and tumor edge regions. GMM preferred to classify tumor edges into three categories which are discrete near the tumor core, and msiPL showed a more discrete distribution in non-tumor sites, especially the region identified as the surrounding vessels **(Fig. 3b)**. We ran SmartGate with different hyperparameters and found that the surrounding blood vessels structure always could be identified, demonstrating its robustness (**Fig. S3**). We also visualized the identified structures and their marker ions’ spatial distribution identified by SmartGate, which showed very clear patterns **(Fig. 3c, d, e)**.

**Fig. 3.**
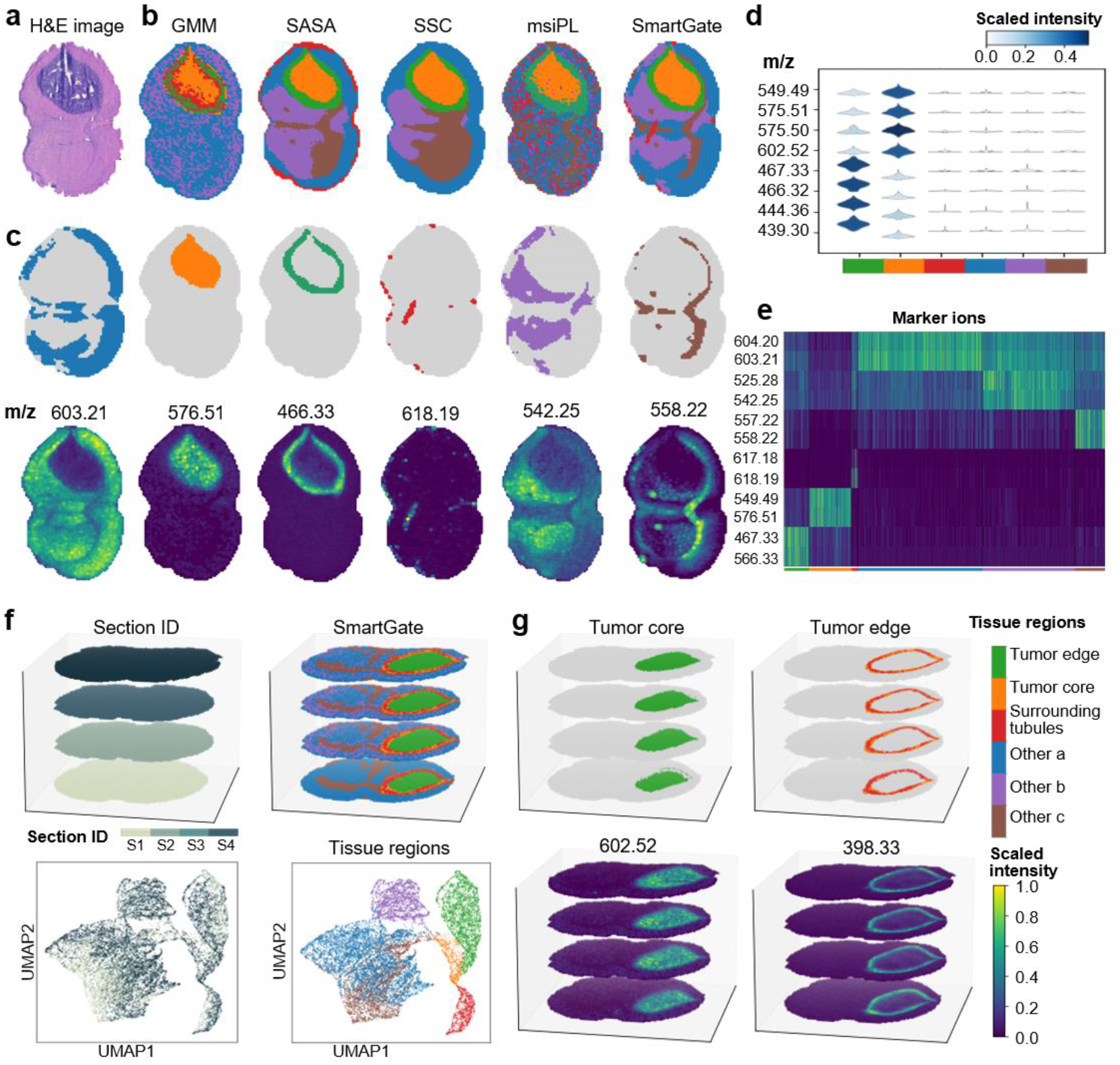
SmartGate reveals the different metabolic patterns of tumor core and tumor edge regions. **a** The H&E images of glioblastoma. **b** Spatial structures identified by GMM, SASA, SSC, msiPL, and SmartGate respectively. **c** Visualization of each structure identified by SmartGate and their marker ions distribution. **d** Violin plot of the marker ions of the tumor core and edge structures identified by SmartGate. **e** Top marker ions determined by differential analysis of each structure identified by SmartGate. **f** Visualization of four tissue sections about the 3D dataset of glioblastoma and the spatial structures identified by SmartGate and the UMAP plot about the section and the spatial structures respectively. **g** The 3D tumor core and edge structures identified by SmartGate and their marker ions’ spatial distribution.

In addition, SmartGate was capable of achieving spatial segmentation of consecutive tissue sections **(Fig. 3f)**, and could well reveal the tumor edge and tumor core regions in the 3D space **(Fig. 3g)**, and the marker ion of the tumor edge (m/z=398.33) through differential analysis was identified as 9-Hexadecenoylcarnitine or trans-Hexadec-2-enoyl carnitine, which was consistent with a recent study [25].

After automatic peak picking, SmartGate obtained 2,719 m/z features and used mclust for the final clustering and differential analysis. We also compared the performance of SmartGate, fMerge, and mclust in terms of the hybrid index and LISI index respectively **(Fig. S4)**, and showed that SmartGate and mclust had similar performance in terms of the LISI index, while fMerge was more discrete than SmartGate and mclust in terms of the distribution of certain categories in the non-tumor part with a different number of clusters. We also tested the effect of selecting different numbers of m/z features on the downstream spatial segmentation **(Fig. S5)** and demonstrated that the automatic peak picking was robust, and the results remained consistent when the number of clusters was set to six.

### SmartGate detects the laminar organization of mouse kidney in both 2D and 3D datasets

We applied SmartGate to the mouse kidney datasets consisting of 95,000 pixels and 140 peaks per pixel with some preprocessing [26] (**Table 1)**. Compared to the mouse tissue structures labeled manually **(Fig. 4a)**, GMM and msiPL divided some known regions like the inner medulla into two parts due to the potential deficiency of spatial information **(Fig. 4b)**. SSC revealed more reasonably segments, but failed to detect some small tissue areas such as lymphatic vessels. Fortunately, SmartGate could extract several parts of lymphatic vessels that are consistent with manual annotations (**Fig. 4c)**, which has also been confirmed by the spatial distribution of the marker ion (m/z 263.04). We also ran a set of hyperparameters of SmartGate (**Fig. S7**) which clearly showed its robustness to identify the lymphatic vessels of the mouse kidney. However, GMM, SASA, SSC, and msiPL could not segment lymphatic vessels like SmartGate even with different numbers of clusters which demonstrated that SmartGate could perform spatial clustering for tissue segmentation more accurately (**Fig. S6**).

**Fig. 4.**
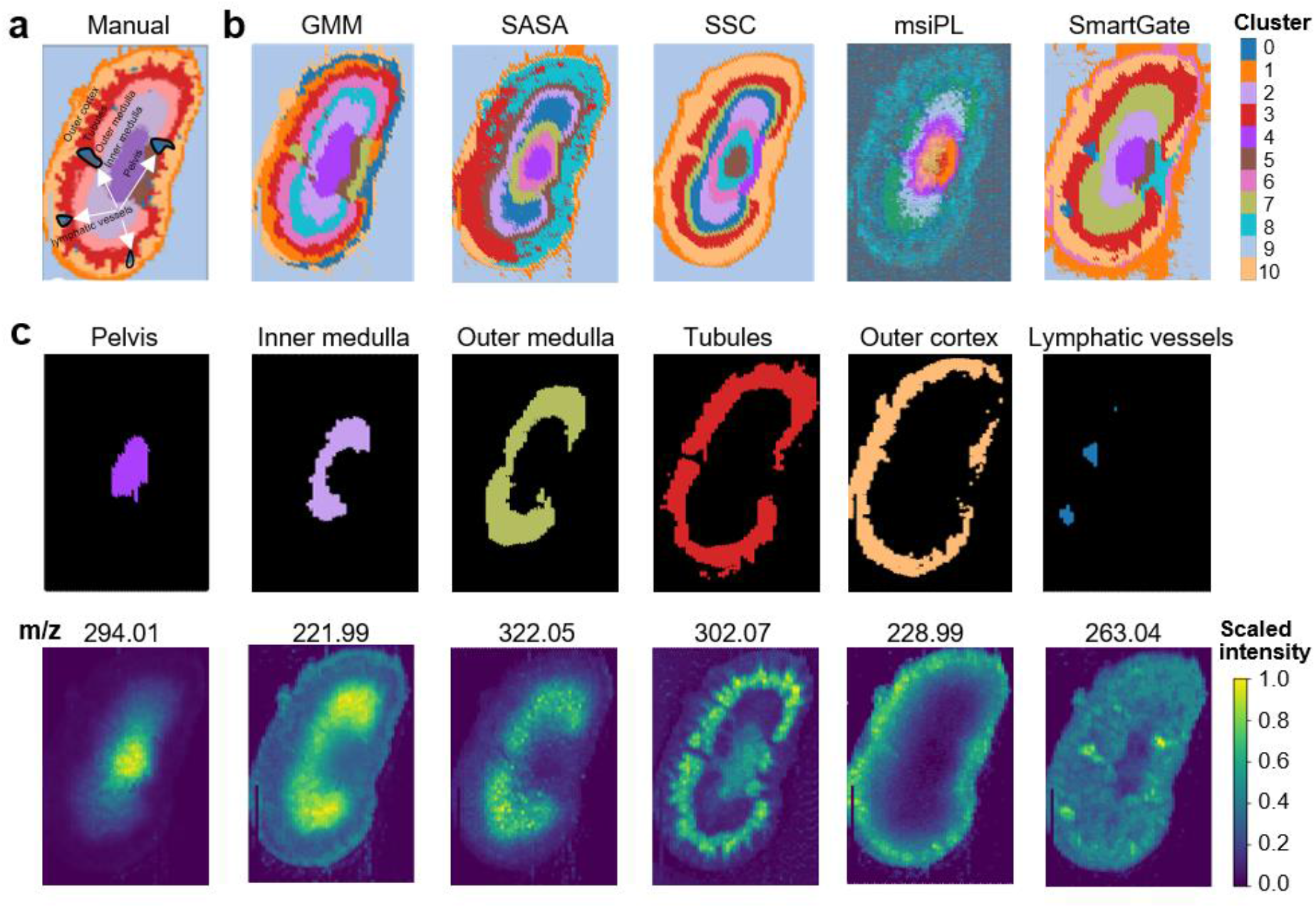
SmartGate detects the different tissue structures of the mouse kidney profiled by nano-DESI. **a** The manual annotation of known spatial structures of the mouse kidney. **b** The spatial structures identified by GMM, SASA, SSC, msiPL, and SmartGate in the mouse kidney dataset respectively. **c** Visualization of each structure identified by SmartGate and its corresponding marker ion spatial distribution.

To gain more understanding of the organization of the mouse kidney, we also used another mouse kidney dataset **(Table 1)** [27]. This dataset contains 75 consecutive sections with a total of 1,362,830 pixels and 7,680 m/z features for each. We first applied SmartGate to individual sections of data which has 18,536 pixels. SmartGate could identify the main tissue structure of the mouse kidney **(Figs. 5a, b)** and divide the renal cortex into three categories (clusters 1, 7, 9), and renal medulla into two categories (clusters 2, 3), which is related to the complex physiological environment within this tissue. Other methods like GMM don’t employ spatial information and thus lead to very discrete results. SASA and SSC extracted some mouse kidney tissues, but SmartGate and msiPL could capture even more accurate segmentation **(Fig. 5a, b)**. We selected six consecutive tissue sections to form a 3D mouse kidney data with a total of 108,475 pixels **(Fig. 5c)**. SmartGate obtained very promising performance compared to that on the 2D data **(Fig. 5d)**, and identified the main labeled tissue structures **(Fig. 5e)**.

**Fig. 5.**
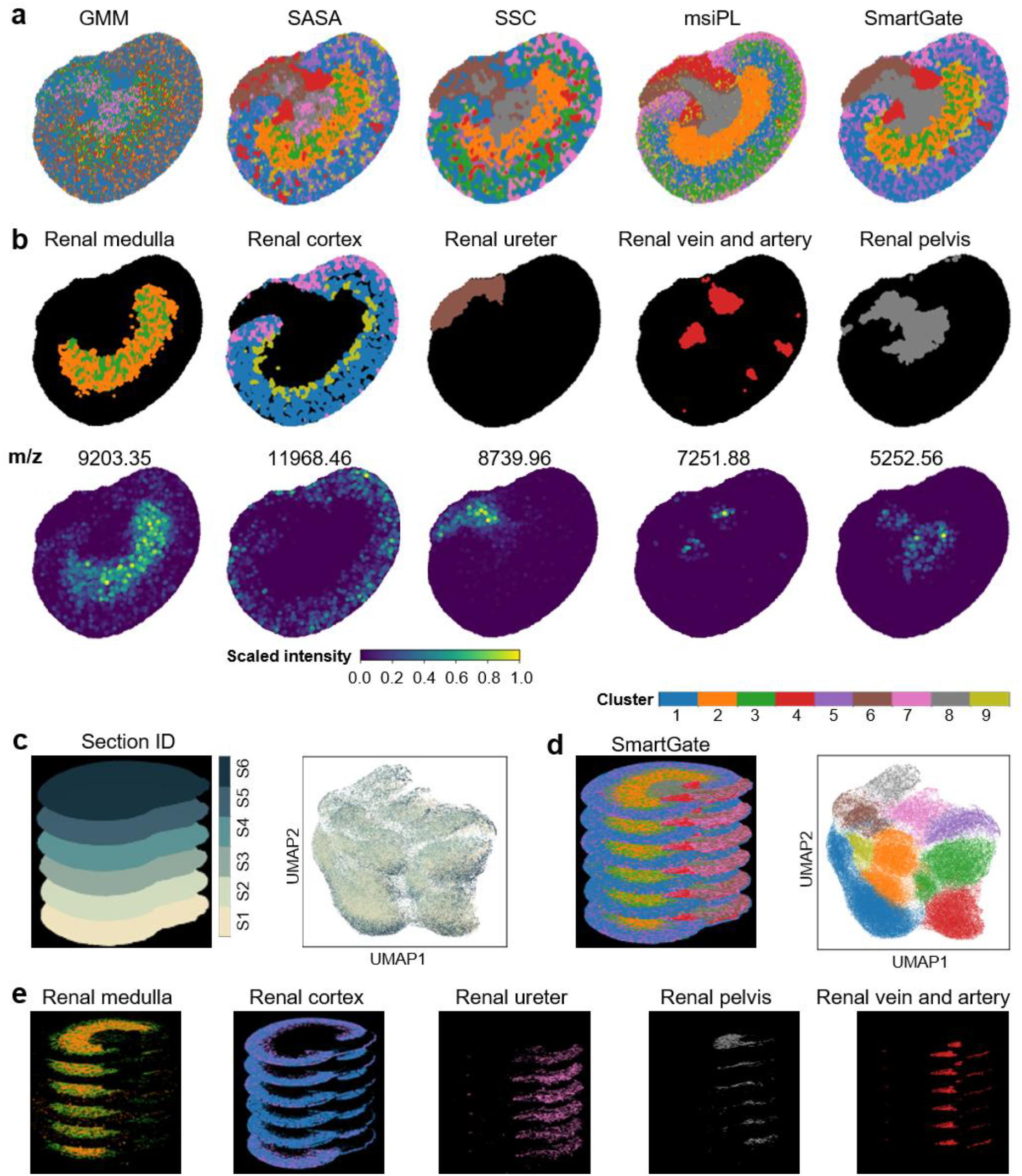
SmartGate identifies the major structures of the mouse kidney profiled by MALDI. **a** The spatial patterns about the mouse kidney identified by GMM, SASA, SSC, msiPL, and SmartGate respectively. **b** The major structures identified by SmartGate and their marker ion spatial distribution. **c** Six consecutive tissue sections of the mouse kidney and the UMAP plot about sections. **d** The 3D Spatial structures identified by SmartGate and the UMAP plot of the spatial structures identified by SmartGate. **e** The major structures of 3D dataset identified by SmartGate.

### SmartGate detects more elaborate spatial structures in the human liver dataset with subcellular resolution

We applied SmartGate to the human liver data collected from liver cancer patients with a fibrotic niche at the subcellular resolution level (**Fig. 6a**) [6]. When we divided all the pixels into six clusters, the metabolic characteristics of each cluster were separated (**Figs. S8a, b, c**). We compared the pixel clusters obtained by SEAM based on only pixels of cellular nuclei (**Figs. S8d, c, e**), and observed that the spatial pattern of each cluster matched closely. Compared with GMM, SASA, SSC, and msiPL **(Figs. 6b, S9)**, SmartGate deciphered the tissue structures more clearly with the clear spatial distribution. GMM and SSC seemed to determine a large cluster without clear spatial patterns. Based on the UMAP plots of the clusters, SmartGate distinguished the spectra of pixels clearly (**Fig. S9b**). Given that the pixel size of subcellular data was much smaller than the size of a cell, we could assume that adjacent pixels were likely to belong to the same cell type. The mean LISIs of each cluster detected by SmartGate were relatively lower than those of other methods. The clusters of SASA were not concentrated as those of SmartGate **(Figs. 6d, S9)**.

**Fig. 6.**
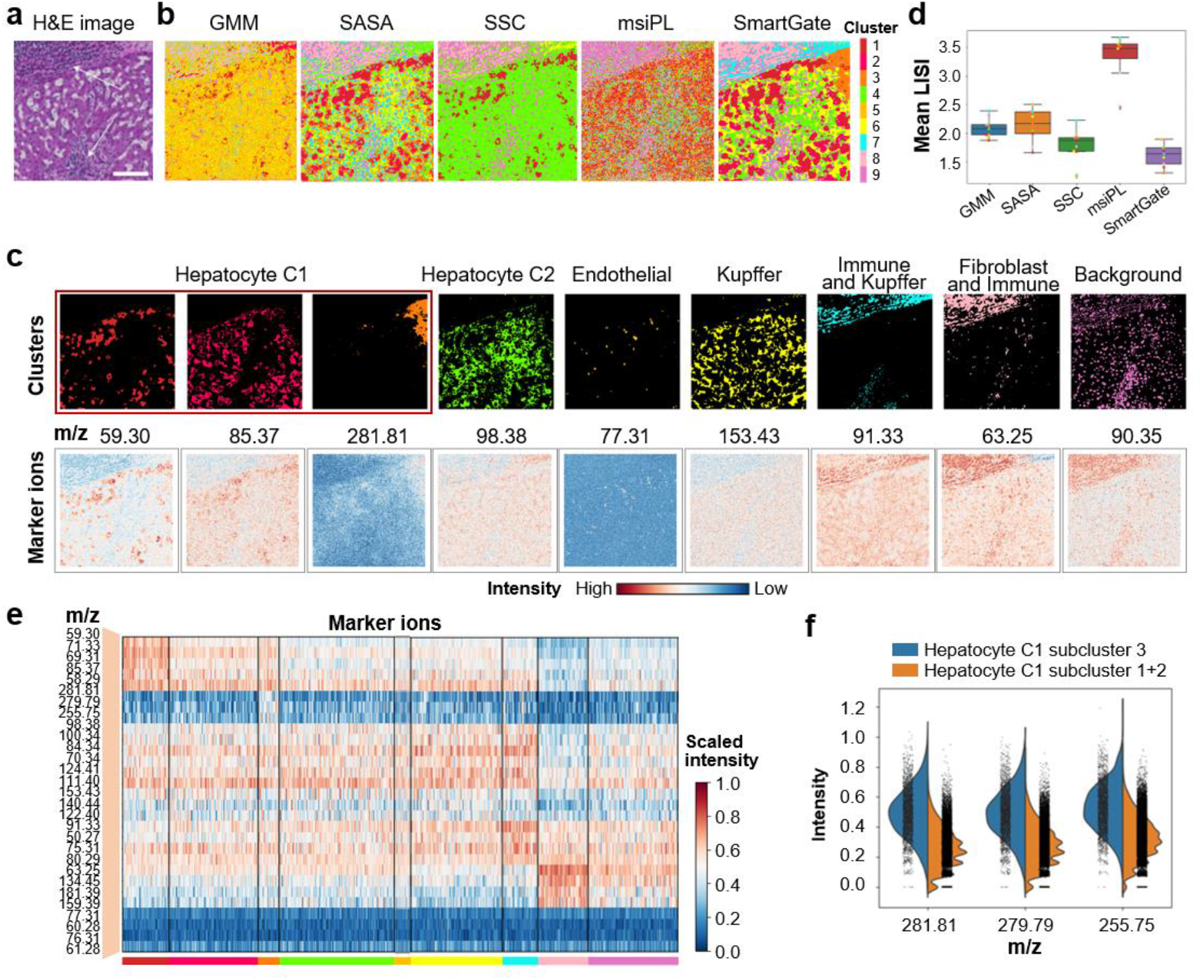
SmartGate detects more elaborate spatial structures of the human liver data. **a** The H&E image. **b** The spatial structures identified by GMM, SASA, SSC, msiPL, and SmartGate respectively. **c** Visualization of each tissue structure identified by SmartGate and its marker ion spatial distribution. One new hepatocyte subpopulation is discovered. Hepa, hepatocyte. **d** Evaluation of clustering performance of these five methods. **e** Top marker ions determined by differential analysis of each structure identified by SmartGate. **f** Comparison of metabolic expression characteristics between the hepatocyte C1 subcluster 1 and the combination of hepatocyte C1 subclusters 2 and 3.

When dividing it into more classes, we could discover more elaborate spatial patterns from the human liver. We annotated the clusters by their marker ions defined by SEAM. These clusters deciphered by SmartGate showed different spatial distributions (**Fig. 6c**) and metabolic characteristics (**Fig. 6e**). We could find the marker ions whose spatial distributions were similar to these clusters, which confirmed the accuracy of segmentation. Hepatocyte C1, which included clusters 1, 2, and 3, was colocalized with m/z 59.3, and cluster 3 was a new hepatocyte subpopulation since its spatial pattern was unique. The new hepatocyte subtype was marked by m/z 281.81, 279.79, and 255.75 (**Fig. 6f**), which were annotated as glycerophospholipid [13]. Compared with other hepatocytes C1, this new cluster was more enriched with these marker ions. Cluster 4 marked by m/z 98.38 was annotated as hepatocyte C2. Most pixels identified by SmartGate belonged to hepatocytes, which was consistent with the fact that 80% of liver cells are hepatocytes [28]. Similar to the results of SEAM, hepatocyte C1 is mainly distributed near the fibrotic niche, while immune and fibroblast cells colocalized with the fibrotic niche. We also adjusted the hyperparameters of SmartGate, the Louvain resolution, and the neighborhood number for the spatial neighbor graph, which showed that the segmentation results were robust (**Figs. S11**).

### SmartGate deciphers the central vein (CV)-centered spatial structures in the mouse liver data

The hepatic lobule is a hexagonal structure with a central vein in the center [29] (**Fig. 7a**). Compared with four methods, SmartGate captured the CV-centered region (red), the little-slice region (yellow), the region spreading from center to the distance (blue) more clearly (**Figs. 7b, S11a**). However, GMM could not detect the CV-centered region and SSC inappropriately split the central vein into two parts. The UMAP plots demonstrated that SmartGate could separate the clusters and the new cluster 8 did have different metabolic characteristics from the other ones (**Fig. S11b**). Moreover, the mean LISIs of clusters identified by GMM was the smallest, but that of clusters identified by SmartGate were more concentrated (**Figs. 7d, S11c**). Note the vastly varied sizes of datasets would severely influence LISI values. The cluster size of GMM was very uneven, resulting in a single cluster dominating most of the neighborhood and some clusters had low LISI values. SmartGate accurately identified the central vein region (pink), while other methods identified some scattered central veins unreasonably.

**Fig. 7.**
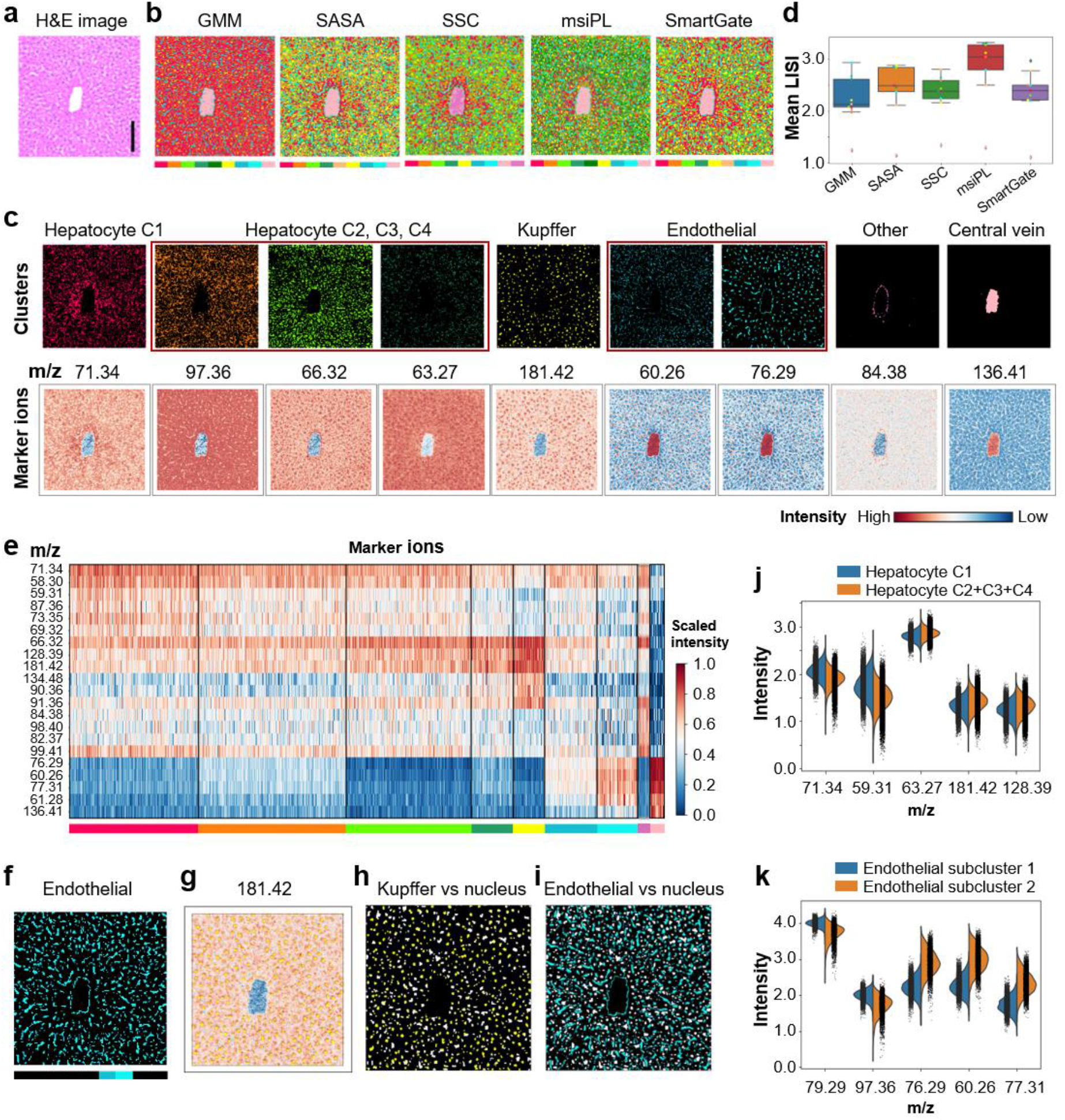
SmartGate deciphers spatial structures of the mouse liver data. **a** The H&E image. **b** Spatial structures identified by GMM, SASA, SSC, msiPL, and SmartGate respectively. **c** Visualization of each tissue structure identified by SmartGate and their marker ion distribution. A new cluster 8 is discovered. **d** Evaluation of clustering performance of these five methods. **e** Top marker ions determined by differential analysis of each structure identified by SmartGate. **f** The spatial structure of endothelial cells. **g** Overlapping of the spatial distribution of Kupffer and its marker ion m/z 181.42. **h** Overlapping of Kupffer cells with cellular nuclei detected by SEAM. **i** Overlapping of endothelial cells with cellular nuclei detected by SEAM. **j** Comparisons of the marker ion intensity between the hepatocyte C1 and the combination of hepatocyte C2, C3, C4. **k** Comparison of the marker ion intensity between the endothelial subclusters 1 and 2.

We labeled the nine clusters according to marker ions of the cell types (**Figs. 7c, e**). The spatial distribution of clusters was consistent with the structure of the mouse hepatic lobule. Hepatocyte C1 was marked by m/z 71.34, and the mixture of hepatocyte C2, C3, and C4 was marked by m/z 97.36, 66.32, and 63.27. Hepatocyte C1 appeared to be spatially enriched around CV and cluster 4 is distally distributed around CV. We observed that the metabolic expression of m/z 71.34, and 59.31 for hepatocyte C1 was higher than those for others, and the expression of m/z 63.27, 181.42, and 128.39 for hepatocyte C2, C3, C4 were higher than those for hepatocyte C1 (**Fig. 7j**). Kupffer cells, the tissue macrophages, were highly colocalized with m/z 181.42 (**Fig. 7g**) and showed the shape of small fragments since macrophages are large. We noticed that Kupffer cells and nuclear cells detected by SEAM (**Fig. 7h**) were highly overlapped which was consistent that their marker ions m/z 90.36, and 134.48 were nuclear markers [13]. Clusters 6 and 7 annotated as endothelial cells showed clear spatial distribution spreading from CV (**Fig. 7f**). Endothelial cells crossed the nuclear region detected by SEAM and were like a line to segment cells with the fact that sinusoidal endothelial was one of compositions of hepatic sinusoids which were along the portal to central vein in the lobules [30] (**Fig. 7i**). The subtypes of endothelial cells also expressed some special ions differently (**Fig. 7k**). We found a new cluster marked by m/z 84.38 and its specific spatial pattern which distributed around CV. The segmentation results by SmartGate with several hyperparameter settings showed no big difference (**Figs. S12**).

## Discussion

Spatial segmentation is a typical method to explore the characteristics of the IMS data and is crucial to understand the tissue structure and metabolic patterns in organisms. SmartGate makes full use of the spatial similarity of adjacent pixels to improve the accuracy of spatial segmentation. SmartGate automatically selects m/z features with potential biological relevance which provided more meaningful embedding features for downstream clustering characterization. SmartGate can be applied to diverse IMS data with different technologies and resolutions.

For low-resolution data, SmartGate can identify substructures in the porcine embryonic brain and different metabolic patterns in tumor edge and tumor core regions, and provide more accurate mouse kidney tissue structures. For datasets with subcellular resolution, SmartGate can leverage all pixel information to segment human liver and mouse liver correctly compared with SEAM only using pixels in cellular nuclei, and discover some new clusters with specific spatial patterns. Moreover, SmartGate can also select potentially more biologically meaningful m/z signatures which would help us identify biomarkers, especially certain disease-specific metabolites. At last, SmartGate can efficiently combine multiple adjacent sections and identify their spatial structures by eliminating the batch effects in 3D IMS data.

In general, preprocessing can reduce the effects of unnecessary noise. As reported by Abdelmoula et al., peak picking can affect the final clustering result and in turn affect the biological interpretation [16]. SmartGate can pick peaks automatically to some extent to alleviate the bias brought by subjective factors.

SmartGate can find new marker metabolites of each spatial pattern. Current studies pay attention to the specific tissues and diseases, like tumors (**Figs. 3, 6, S13**) [11, 25, 31], but the ions or metabolites of some tissues obtained by differential analysis lack clear biological information, which hinds the debugging and development of SmartGate to a certain extent. In the future, we hope to further increase the interpretability of SmartGate with the help of other omics such as single-cell metabolomics and spatial transcriptomics for multi-modal omics integration [32] and can use spectrometry to predict molecular formulas and thus identify metabolites.

## Materials and Methods

### Materials

We applied SmartGate to six diverse datasets (**Table 1**). The pig fetus dataset consisting of 4,959 pixels was generated by a Thermo Finnigan LTQ linear ion trap MS -DESI over the range of 150 –1,000u m/z [18]. The spatial resolution for each pixel was 100 μm, and 10,200 peaks were first selected by experts to construct the metabolic matrix. The glioblastoma (GBM) dataset of the patient-derived xenograft (PDX) mouse brain model consisting of 3,570 pixels and 21,241 peaks was profiled by a MALDI FT-ICR IMS platform. Spectra were collected in the range of m/z 100– 3,000u. The resolution of this dataset is 100 μm [25]. The two mouse kidney datasets were generated by nano-DESI and MALDI respectively. The nano-DESI data consists of 95,000 pixels and 140 peaks per pixel. The dataset by MALDI mass spectrometer was in the mass range of m/z 2,000-20,000u and contained 75 consecutive slices with a total of 1,362,830 pixels, and each with 7,680 m/z features. The resolution of both the datasets were 50 μm. The human and mouse liver data were generated by the time-of-flight-secondary ion mass spectrometry (TOF-SIMS) with 0-2000u mass range [6]. These datasets contained 200-500 peaks. These tissue areas were 400×400 μm^2^ and 256×256 pixels with subcellular resolution of 1.5 μm per pixel.

### Data preprocessing

For all datasets, we adopted the total ion count (TIC) normalization to normalize the original data of each pixel by their total intensity over all ions. If *X*_*i*_(*i* = 1, …, *N*) ∈ *R*^*p*^ is the spectrum features of the *i*-th pixel, TIC-normalization is defined as follows,

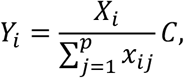

where *C* is an empirical constant. For the multicellular resolution data, *C* is set to the default value 1. For the subcellular resolution data, *C* is set to their mean intensity accordingly (i.e., 200 for the human liver data and 220 for the mouse liver data).

### Graph attention autoencoder

Graph attention autoencoder [33] is an unsupervised learning neural network architecture which reconstructs the features of nodes by learning the relationships of nodes in the graph structure using autoencoder. The encoding layer generates new features of each node by the features of their neighbors, using self-attention mechanism to achieve parameter sharing among nodes. The decoding layer uses the same number of layers as the encoding layer to invert the encoding process for the purpose of reconstructing node features. The low-dimensional embedding feature vectors of the hidden layer in the auto-encoder are obtained by training which are used for downstream analysis. STAGATE adapts this strategy for deciphering spatial transcriptomics to extract low-dimensional features for further clustering and applications. SmartGate adopts this strategy with the auto-encoder consisting of 215-10-215 neurons for the multicellular resolution data and 512-30-512 neurons for the subcellular resolution data respectively.

### Clustering algorithm

SmartGate takes the low-dimensional representation of each pixel as the input of the clustering algorithm. Louvain algorithm implements a network community division strategy [23] and mclust implements a Gaussian finite mixture density model using R [34]. For datasets with different biological backgrounds, we specify the number of clusters in advance.

### Details of automated peak picking

We take the original IMS data as input for the first iteration, and use the Graph attention autoencoder to reduce the dimensionality to obtain a low-dimensional embedding vector, and then input the low-dimensional vector into mclust to obtain the initial spatial structure, in order to reduce the influence of noise in the original data, we suggest a small number of clusters after the first iteration. In this paper, we set the number of categories to four, and then use the difference analysis to calculate the specific ions for each category. We select the first 800 ions for each category, and input the final m/z features into the autoencoder again to obtain the final embedding vector. Alternatively, multiple iterations can be performed, e.g., selecting the top 1600 ions for each category and then iterating here **(Fig. S5)**.

### fMerge

fMerge is the process of dimensionality reduction of the original data by selecting a sliding window for the original mass spectrometry results and calculating the mean value for all peaks in the window.

### Differential peak learning

To find the marker ions for each cluster or cell type, the differential peak learning is analyzed on the metabolic features by rank_genes_group from SCANPY. It tests the target cluster and the rest clusters by student’s t-test and rank the p-value to find the most different expressed ions and the Wilcoxon test algorithm, which is commonly used as a nonparametric test, is also able to identify more m/z features with specificity. When there are some clusters sharing some ions, we need to select the top and specific ions for each cluster.

### Comparisons of other segmentation algorithms

#### LISI Index

LISI is proposed to assess mixing and accuracy of clustering results from Harmony [35]. First, LISI finds the neighborhoods of pixels by building Gaussian kernel distribution of the distances of the neighborhoods and fixing the perplexity which can be regarded as the expected number of each pixel’s neighborhoods. Then, LISI computes the expected sampling number of pixels before two pixels belonging to the same cluster are draw, that is the LISI value of a pixel as follows,

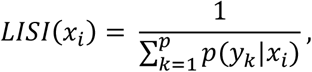

where *x*_*i*_ is the i-th pixel, *y*_*k*_ is the cluster of the neighborhood of the i-th pixel, *p* is the complexity fixed to 8 and 12 for high and low resolution datasets respectively. Here, we calculate the spatial location distance between neighborhoods and compute LISI by Lisi R package. To observe the mixing of each cluster, we calculate the mean LISI for each cluster and draw a boxplot. More intuitively, we draw the kernel density plots of each cluster by Seaborn.

#### Hybrid Index

The formula of Hybrid index is as follows,

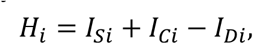

while *I*_*Si*_, *I*_*Ci*_, *I*_*Di*_ are the Silhouette coefficient, Calinski-Harabasz score and Davies-Bouldin score of *i*-th method respectively.

### Enrichment analysis of the clustering results

To compare the segmentation results with SEAM which generates the data, the enrichment analysis between SmartGate and SEAM is carried on the nuclear pixels selected from SEAM. We perform Fisher exact test on the 2×2 confusion matrix composed of a certain cell type by Scipy and the remaining cell type of SEAM and a certain cluster and the remaining cluster of SmartGate, and then perform FDR correction on p-value by Statsmodels. We regard P-value < 0.05 as significant enrichment, otherwise it is not significant.

### Datasets

The datasets used in this study are all from published data sets (**Table 1**). The pig fetus dataset is accessible within the R cardinal package (https://www.bioconductor.org/packages/CardinalWorkflows/). The PDX-GBM dataset was collected at (https://www.metabolomicsworkbench.org/search/sitesearch.php?Name=PR001171). The mouse kidney by nano-DESI dataset is available at (https://github.com/hanghu1024/MSI-segmentation). The mouse kidney by MALDI datasets is available at (http://gigadb.org/dataset/100131). The human liver and mouse liver datasets are available at Github (https://github.com/yuanzhiyuan/SEAM/tree/master/SEAM/data/raw_tar).

## Code availability

The SmartGate package is implemented in Python and is available on Github [https://github.com/zhanglabtools/SmartGate].

## Acknowledgements

This work has been supported by the National Natural Science Foundation of China [Nos. 12126605, 61621003 to S.Z.], the National Key Research and Development Program of China [No. 2019YFA0709501 to S.Z.], the Strategic Priority Research Program of the Chinese Academy of Sciences (CAS) [Nos. XDA16021400, XDPB17 to S.Z.], the Key-Area Research and Development of Guangdong Province [No. 2020B1111190001 to S.Z.].

## Author contributions

S.Z. conceived, desinged and supervised the project. K.X., Y. W. and K.D. developed and implemented the SmartGate package. K.X., Y. W., K.D. and S.Z. validated the methods and wrote the manuscript. All authors read and approved the final manuscript.

