## Supplementary Text, Figures and Tables. for "SmartGate is a spatial metabolomics tool for resolving tissue structures"

#### This PDF file includes:

Supplementary Text, Figures and Tables.

### Supplementary Text

#### Four existing methods

**Gaussian mixture model (GMM)** [34] It assumes that all data follows the normal distribution. Different subgroups are learned from the mixed distribution, and the data in each subset follows a normal distribution. To estimate the parameters of the normal distribution of different subsets, GMM uses the maximum expectation (EM) algorithm to estimate the parameters. Assuming that a total of *K* subsets can be divided and each subset follows the normal distribution, the mixture distribution of all data is obtained as:

$$p(x) = \sum_{i=1}^{K} w_i N(x|\mu_i, \sigma_i),$$

where $w_i$ is the weight of each subset, $\mu_i, \sigma_i$ are the Gaussian distribution density function of the $i$th sub-model. We implemented GMM using the scikit-learn package and the hyperparameter is the number of mixture components.

**Spatially aware structure-adaptive (SASA)** [17] It uses spatial-aware structure-adaptive distance to define the distance of spectra $d(S_1, S_2)$ between two pixels $(x_1, y_1), (x_2, y_2)$. The definition is as follows:

$$d(S_1, S_2) = \sum_{-r \le i,j \le r} \vec{\alpha}_{ij}(S_1, S_2) \| S(x_1 + i, y_1 + j) - S(x_2 + i, y_2 + j) \|_2^2,$$

$$\vec{\alpha}_{ij}(S_1, S_2) = \alpha_{ij}(S_1, S_2) \sqrt{\beta_{ij}(S_1)\beta_{ij}(S_2)},$$

where $\vec{\alpha}_{ij}$ is the weight of the neighbors in the SASA distance. $\alpha_{ij}$ follows Gaussian distribution which means that $\alpha_{ij} = \exp\left(\frac{-(i^2+j^2)}{2\sigma^2}\right)$ with $\sigma = \frac{2r+1}{4}$, and *r* is the number of neighbors. $\beta_{ij}(S) = \exp\frac{-\delta_{ij}(x,y)^2}{2\omega^2}$ , where $\omega$ is an empirical parameter and $\delta_{ij}(x,y) = \|s(x+i, y+j) - s(x,y)\|_2$.

The adaptive distance between two spectra needs to use K-means algorithm to cluster finally. SASA was implemented by *spatialKMeans* in the Cardinal package and specified different number of clusters with the 'adaptive' method. For the human and mouse liver dataset, we set the spatial neighborhood radius is 1.3.

**Spatial-shrunken centroids (SSC)** [18] K clusters are segmented by SA or SASA distance and the centroid of each cluster $\bar{X}_i$ $(i = 1 \ldots . K)$ is shrunken to the centroid of all data $\bar{X}$. For the spectra feature $p$ for cluster k, the t-statistical value is

$$t_{kp} = \frac{\bar{X}_{kp} - \overline{X_p}}{\widehat{\tau_p}\sqrt{\frac{1}{N_k} - \frac{1}{\sum_{k=1}^{K} N_k}}},$$

where $\widehat{\tau_p}$ is the estimate value of the within-class standard deviation for the feature $p$. After shrinking the centroid of each cluster, the intensities of the shrunken centroids are:

$$\bar{X}'_{kp} = \bar{X}_p + t'_{kp}\widehat{\tau_p}\sqrt{\frac{1}{N_k} - \frac{1}{\sum_{k=1}^{K} N_k}},$$

where $t'_{kp} = sign(t_{kp})(|t_{kp}| - S), t_+ = t \ if \ t > 0,$ and $t_+ = 0 \ if \ t \le 0$ . $s$ is the shrinkage parameter which usually need to be given according to the dataset. When get the new centroid of each cluster, we need to calculate the discriminant score $D(x_{ij}, \bar{X}'_k)$ using SA or SASA distance and the probability that different pixels belong to the same category

$$\widehat{p_k}(S) = \frac{e^{-\left(\frac{1}{2}\right)D(x_{ij},\bar{X}'_k)}}{\sum_l^K e^{-\left(\frac{1}{2}\right)D(x_{ij},\bar{X}'_l)}}.$$

We implemented this algorithm using *spatialShrunkenCentroids* in the Cardinal package. For the pig fetus dataset and PDX-GBM datasets, we set $s = 10$, the radius is 2 and the weights follow Gaussian distribution. For the mouse kidney, we set $s = 5$, the radius is 2 and the weights follow Gaussian distribution. For the human and mouse liver dataset, we set $s = 0$, the radius is 1.3 and the weight is adaptive.

**msiPL** [15] uses variational autoencoder (VAE) to reduce the dimension of the original dataset and uses GMM to classify the reduced dimensional spectral features. VAE network is an unsupervised dimensionality reduction algorithm including encoder, decoder and the hidden layers which have three fully connected neural network. VAE used batch normalization (BN) to regularize the neural network. We used the default parameters with the number of neurons in the mildest layer as 5.

**Hyperparameter Setting** The hyperparameters of SmartGate are the weight of the self-attention mechanism of cell-type aware $\alpha$ [19], the pre-defined radius $r$, the pre-cluster Louvain resolution (if $\alpha \neq 0$), and the embedding-cluster Louvain resolution or the number component of mclust. For all datasets, we summarize the values for these hyperparameters **(Table S1).**

**SmartGate identifies cancer regions on the human prostate cancer dataset.**

We further tested the performance of spatial segmentation by SmartGate on the human prostate cancer dataset profiled by FT-ICR. The original data has 12,716 pixels and 61,343 peaks per pixel. The resolution is 120 μm. Compared to the location of the cancer manually **(Fig. S13a)**, GMM and msiPL could not segment continuous boundaries in the inner of tissue. All these methods could identify the aero of cancer on the right middle while the aero of cancer on left upper are difficult to segment due to the interference of normal surrounding tissue **(Fig. S13b)**. SmartGate could well reveal the cancer tissue (cluster 3) and another clear tissue structure (cluster 2) which could be identified by all methods (**Fig. S13c**). Besides the artificially labeled cancer sites, there were still areas of cancer elsewhere and the spatial distribution of marker ion also prove the rationality of clustering. All the differential marker ions illustrated the specificity of most major structures by SmartGate (**Fig. S13d**).

#### Supplementary Figures

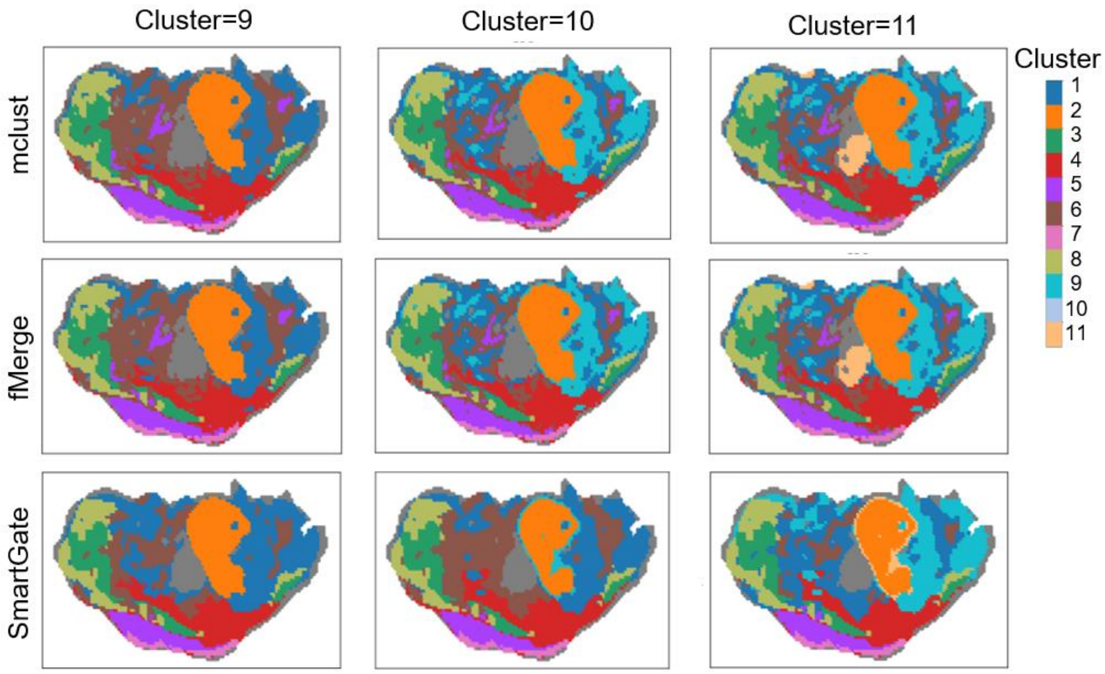


**Fig. S1**. Comparison of spatial structures identified by mclust, fMerge and SmartGate with different number (9, 10, 11) of clusters in the pig fetus dataset respectively.

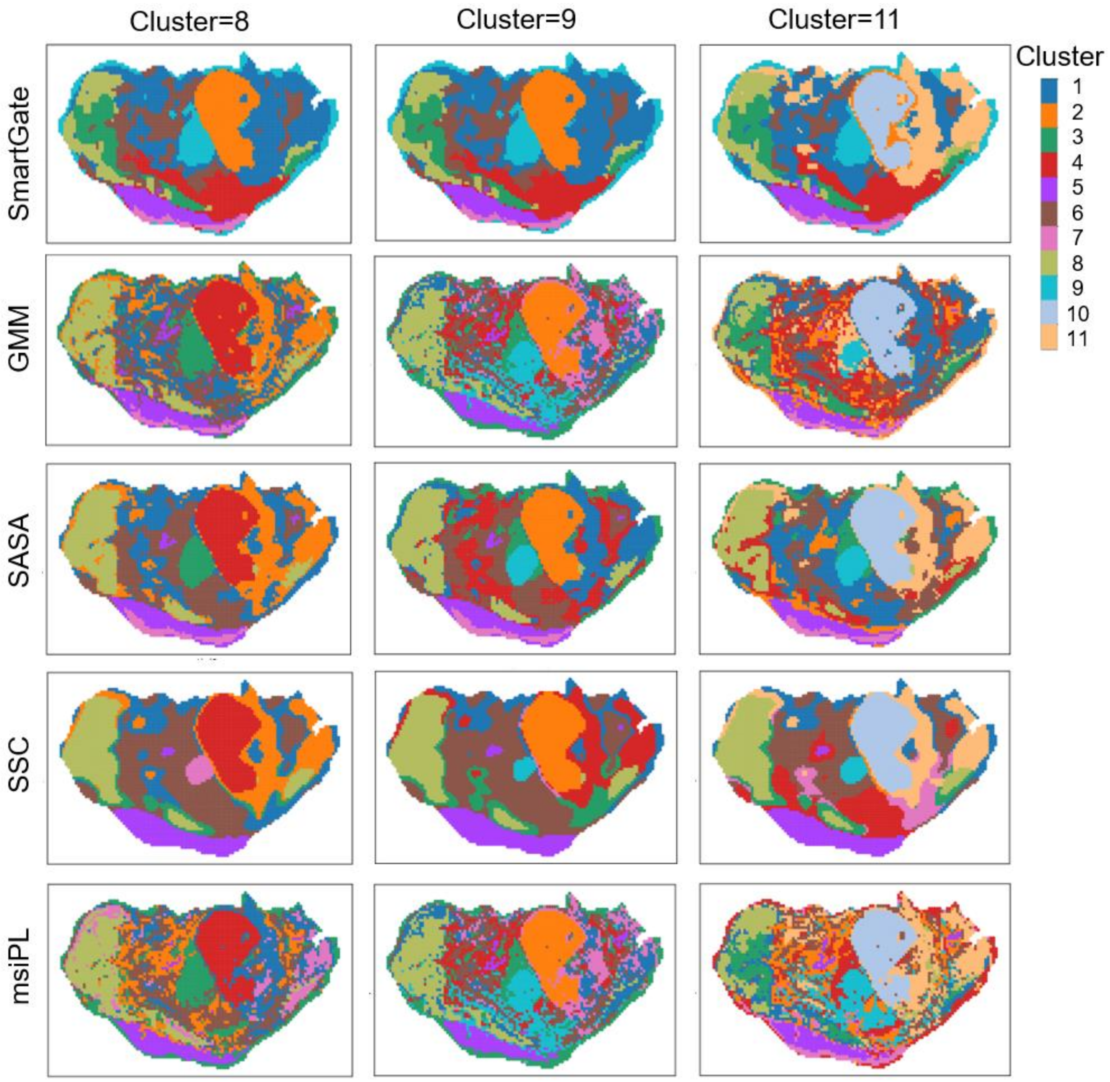


**Fig. S2**. Comparison of spatial structures identified by SmartGate, GMM, SASA, SSC and msiPL with different numbers (8, 9, 11) of clusters in the pig fetus dataset respectively.

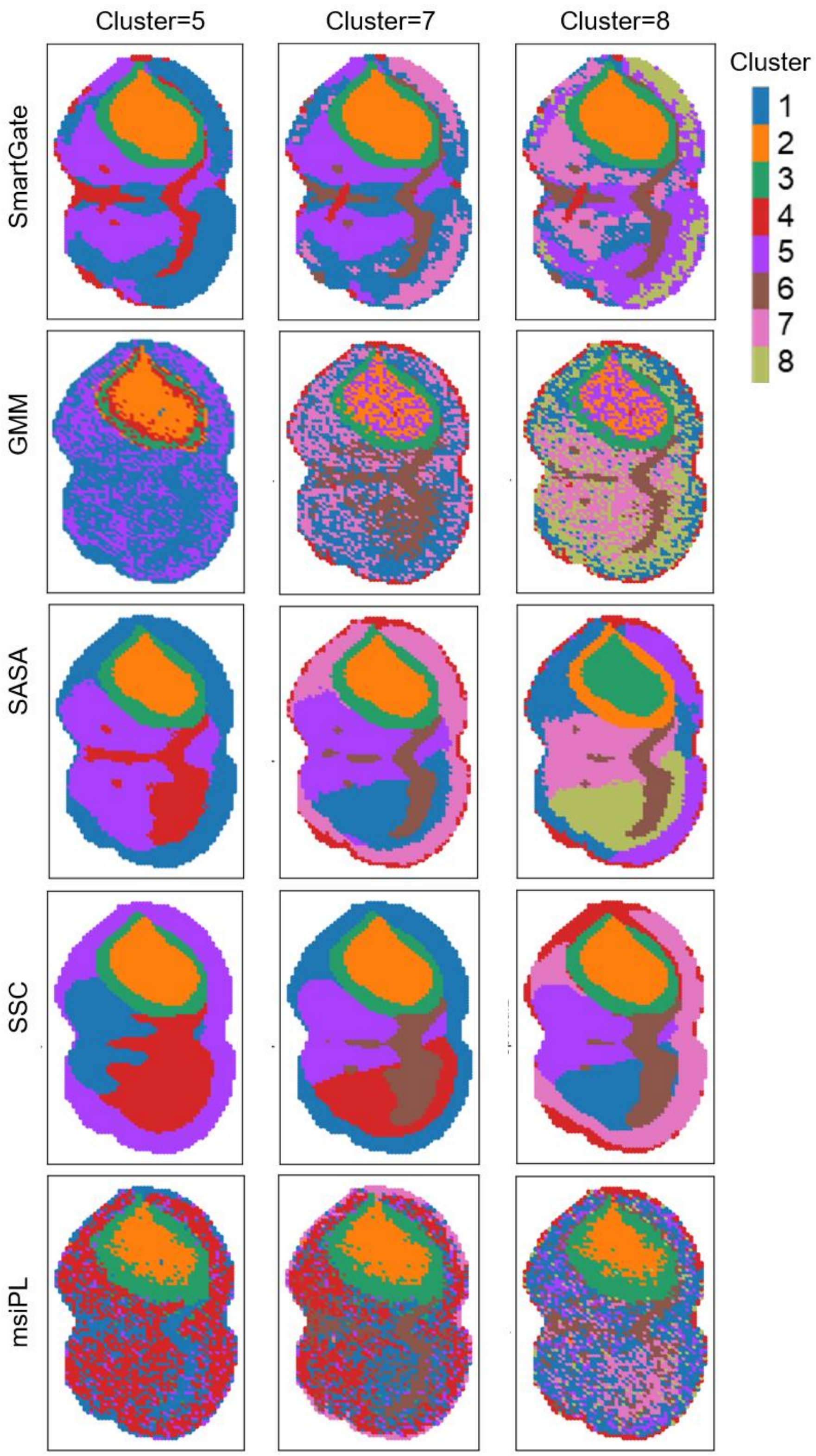


**Fig. S3**: Comparison of the results of spatial segmentation in glioblastoma (GBM) datasets using SmartGate, GMM, SASA, SSC and msiPL with different numbers (5, 7, 8) of clusters in the in the GBM dataset.

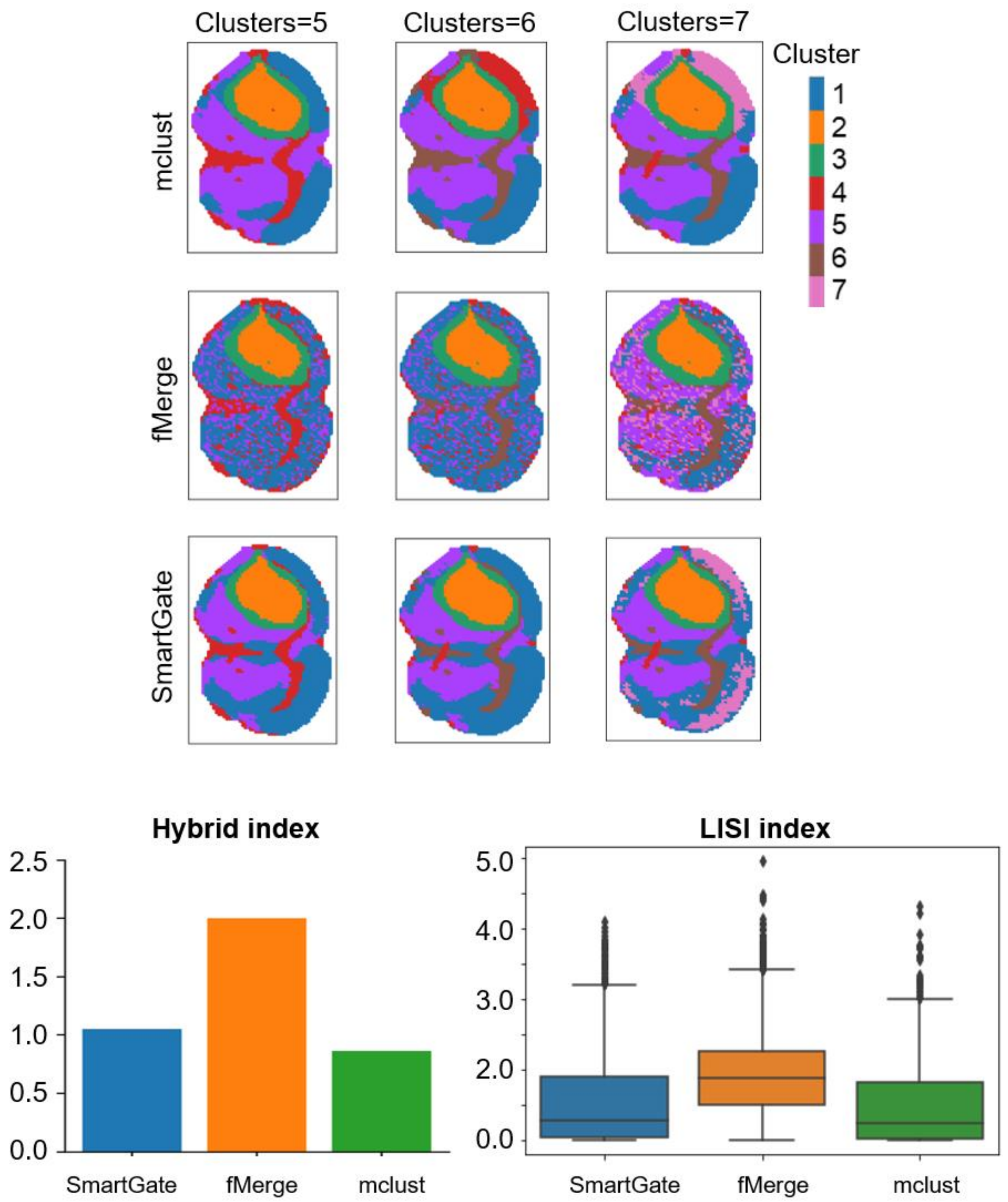


**Fig. S4**. Comparison of spatial structures identified by mclust, fMerge and SmartGate with different number (5, 6, 7) of clusters in the GBM dataset respectively and corresponding comparison in terms of the hybrid index and LISI index respectively.

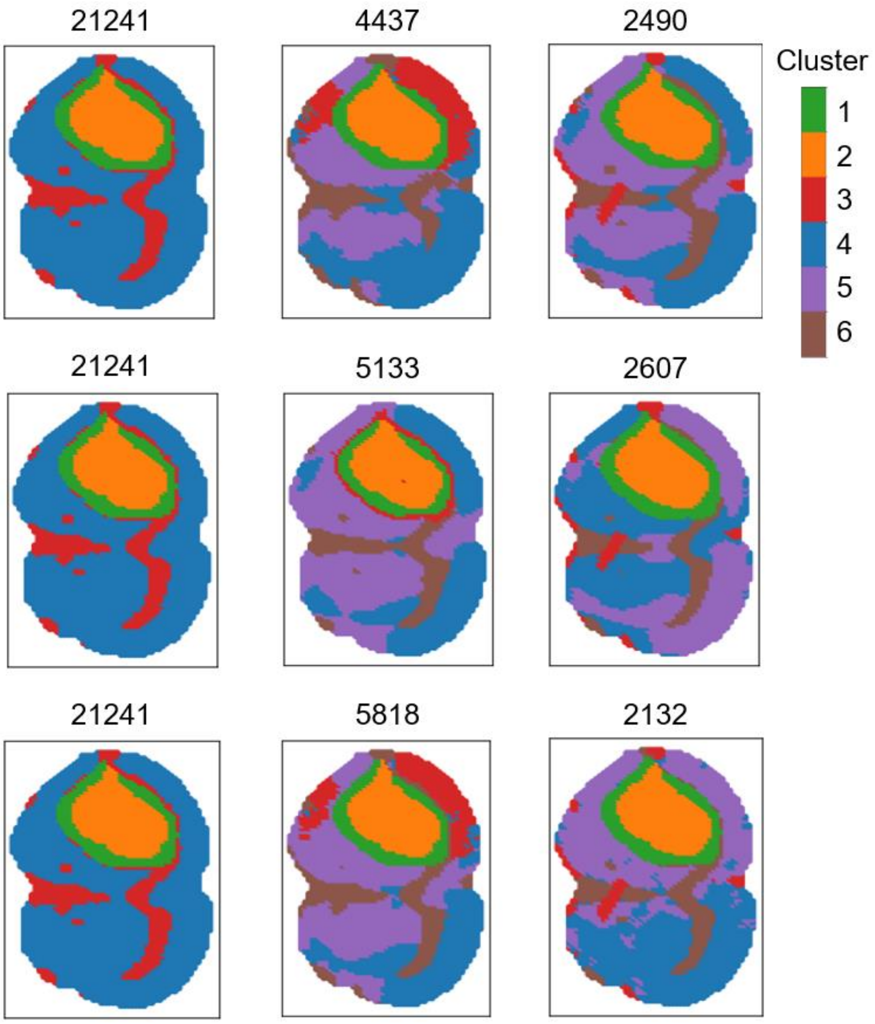


**Fig. S5**. Comparison of spatial structures identified by SmartGate by selecting different numbers of m/z features.

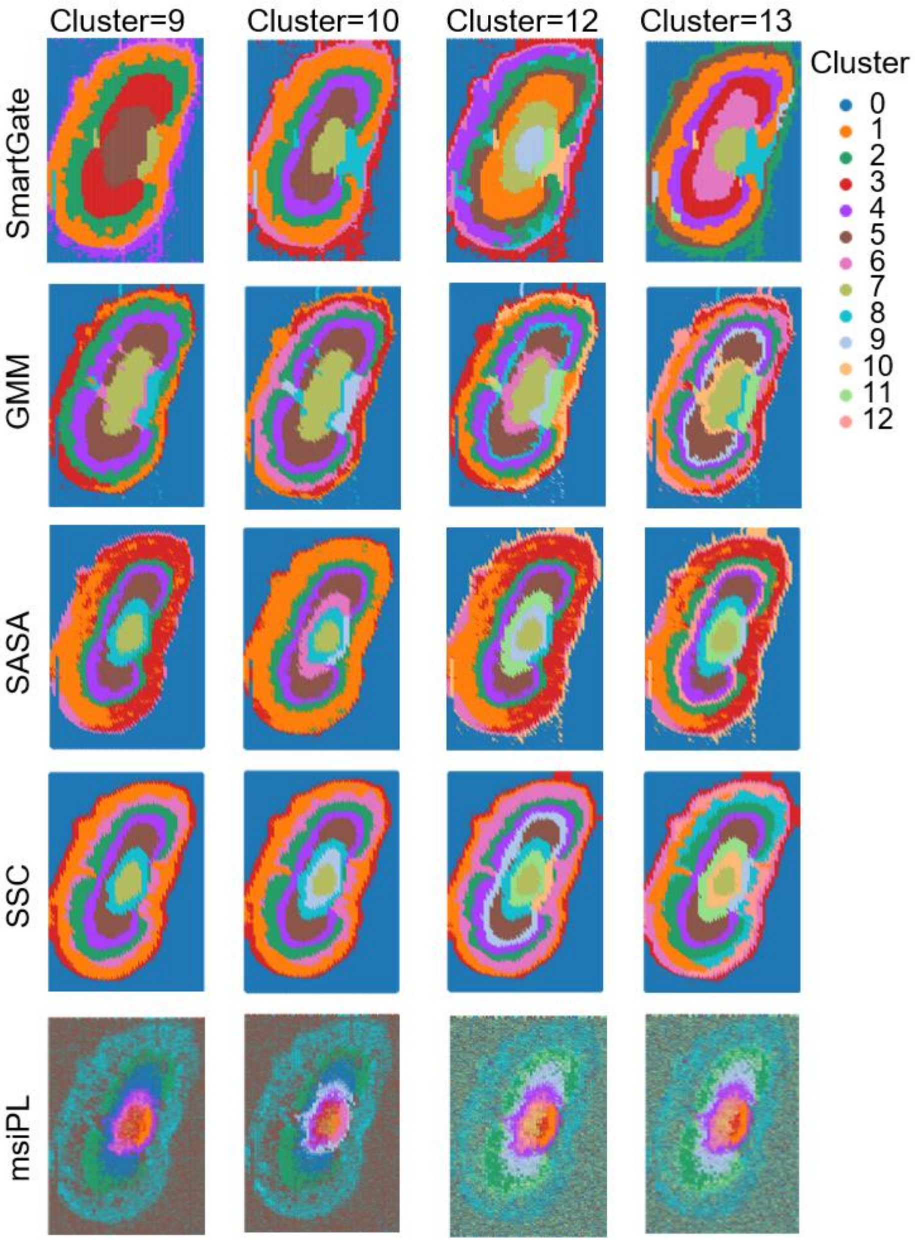


**Fig. S6**. Comparison of spatial structures identified by SmartGate, GMM, SASA, SSC and msiPL with different numbers (9, 10, 12, 13) of clusters in the mouse kidney dataset respectively.

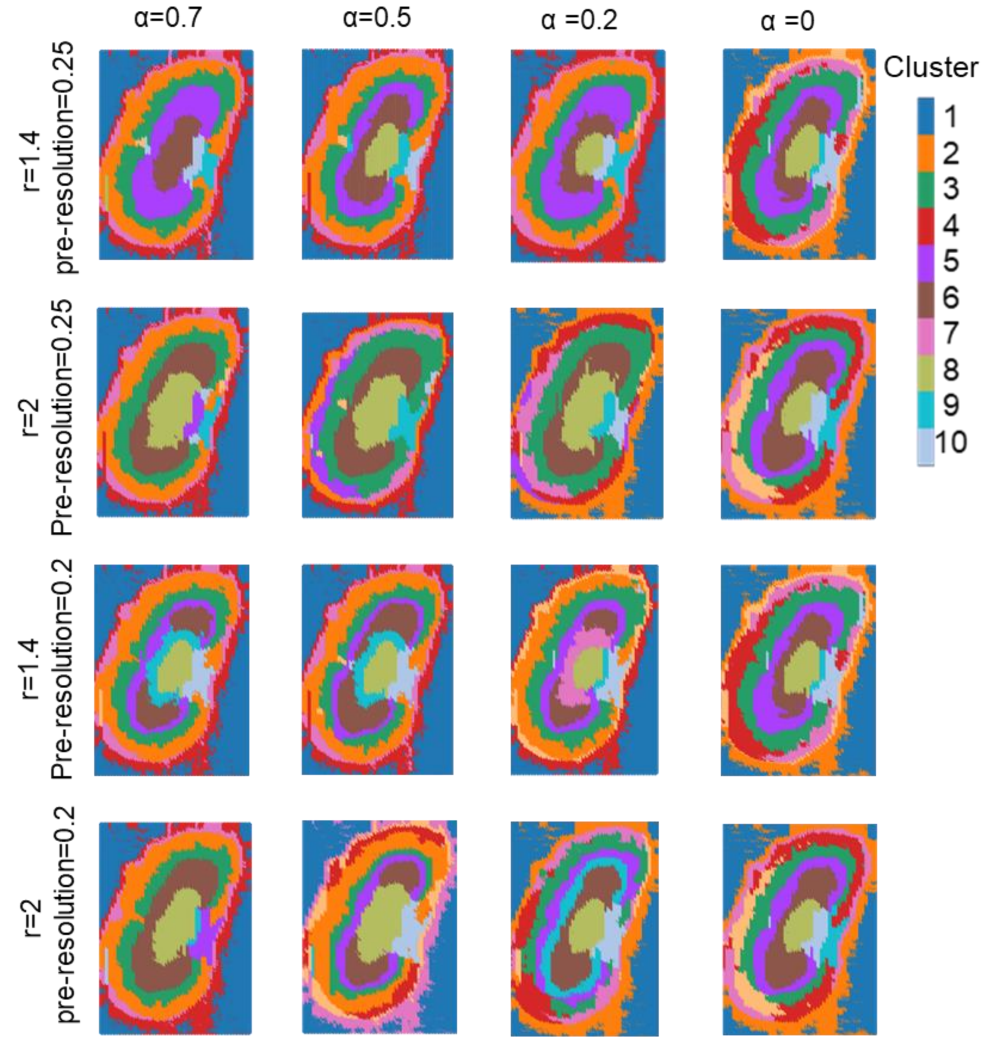


**Fig. S7.** Comparison of spatial structures identified by SmartGate with different hyperparameters ($r$=1.4, 2, pre-resolution(pre-cluster Louvain resolution)=0.2, 0.25 and $\alpha$=0, 0.2, 0.5, 0.7) in the mouse kidney dataset respectively.

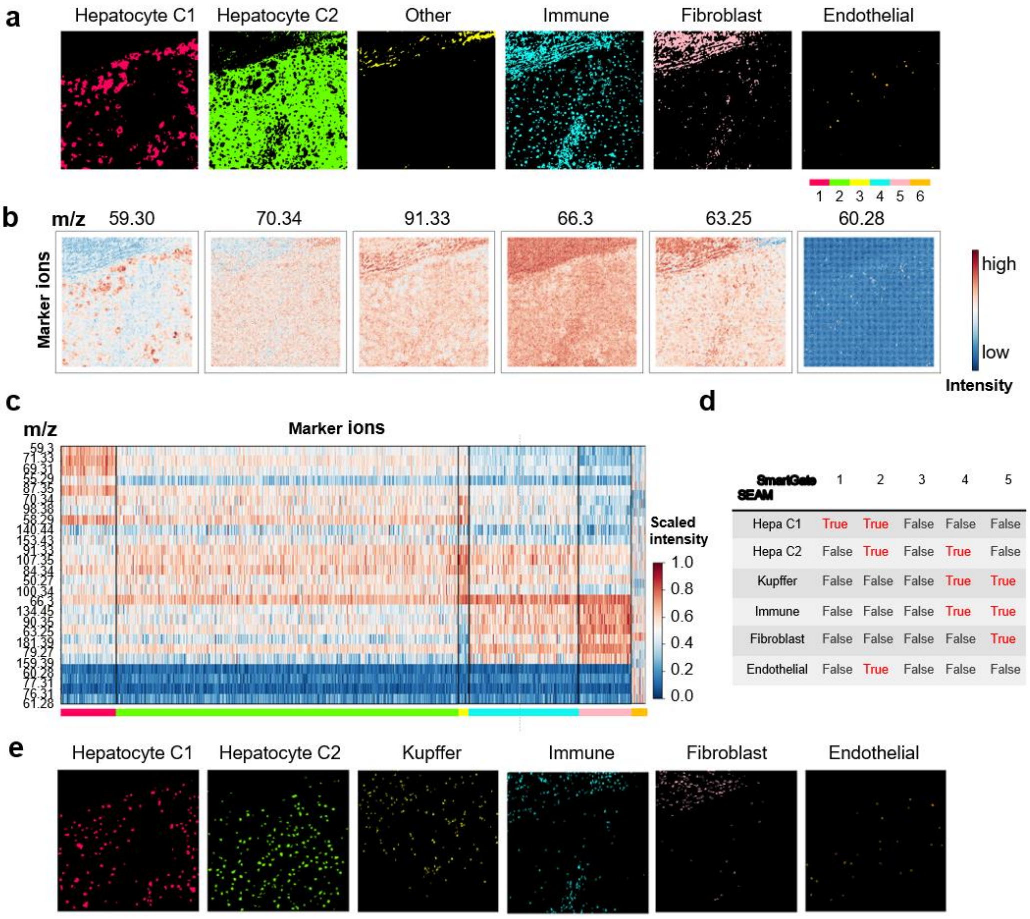


**Fig. S8. SmartGate deciphers relatively consistent spatial structures in human liver data with SEAM. a** Spatial structures identified by SmartGate with the Louvain clustering. **b** Spatial distribution of their marker ions. **c** Top marker ions determined by differential analysis of each structure identified by SmartGate. **d** Overlap analysis between the spatial structures identified by SmartGate and the annotated cell types by SEAM based on the selected nuclei. **e** Spatial single nucleus map of SEAM. The hyperparameters of SmartGate are $\alpha = 0,\ r = 1.3$, and embedding-cluster resolution 0.3.

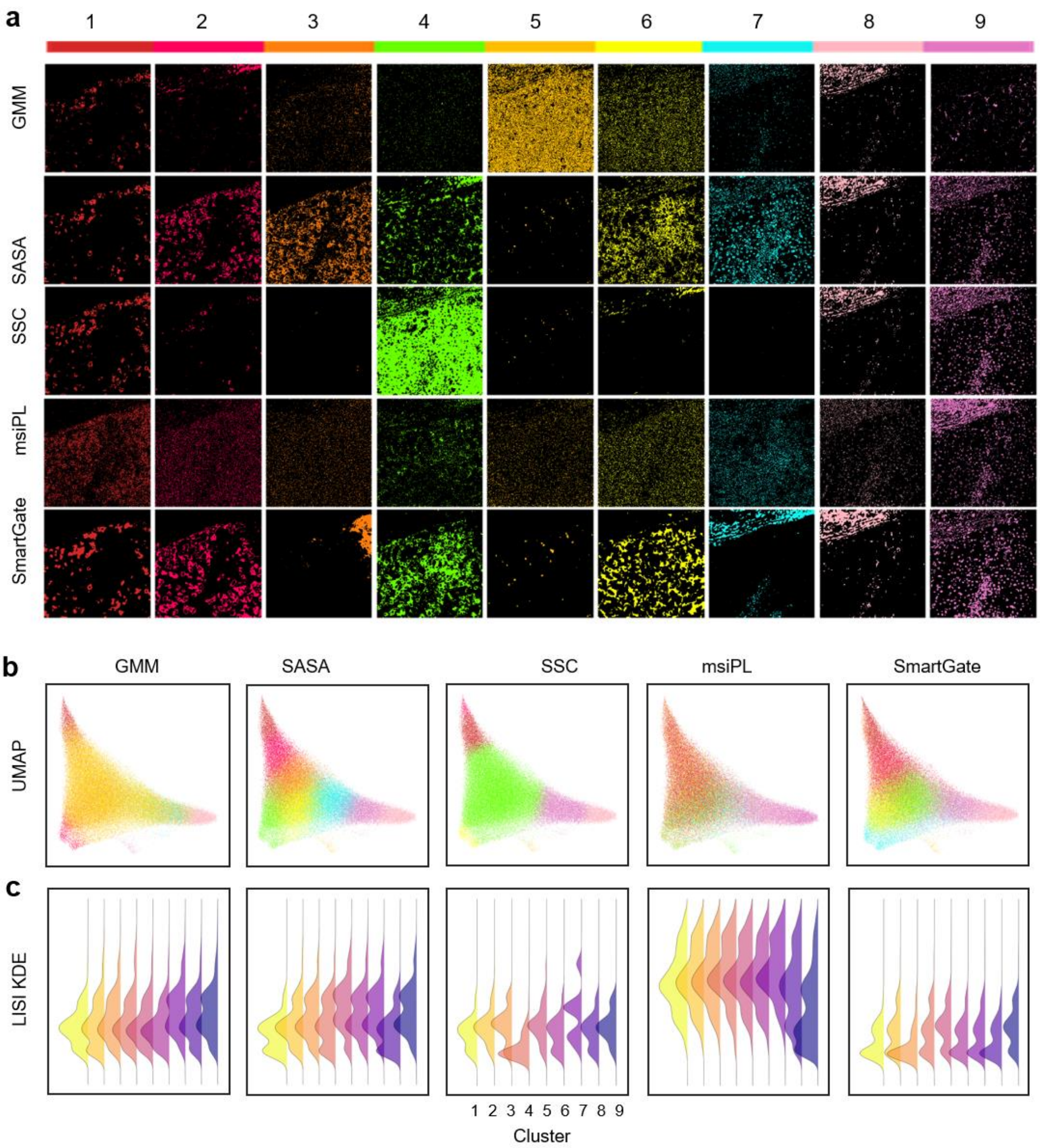


**Fig. S9. Comparisons of the spatial structures by SmartGate and other four methods in the human liver data. a** Each spatial structures identified by GMM, SASA, SSC, msiPL and SmartGate with the Louvain clustering. **b** The UMAP plots of spatial structures by the five methods respectively. **c** The kernel density plot of the LISI values for each structure of the five methods.

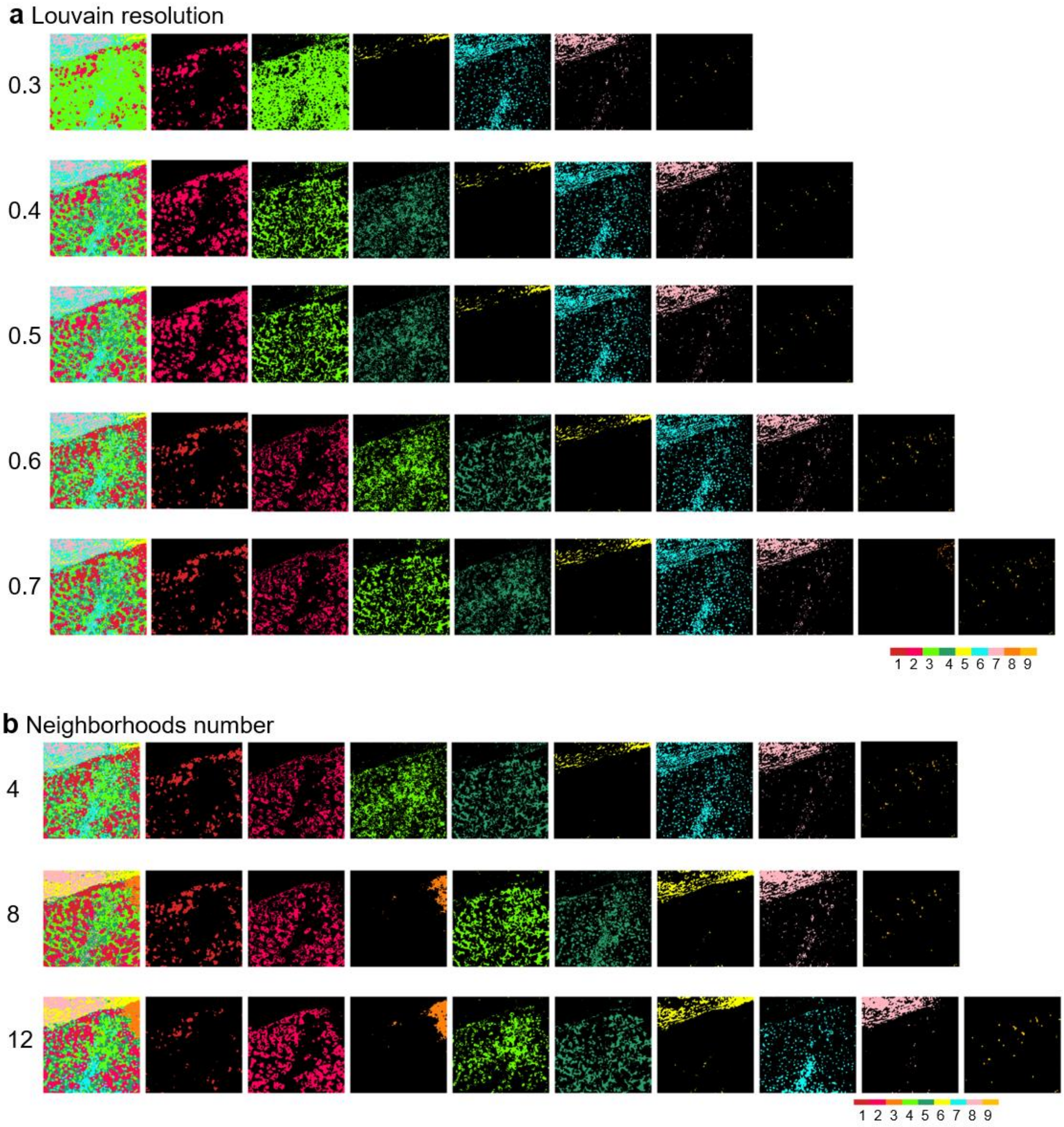


**Fig. S10. Comparison of spatial structures identified by SmartGate with diverse Louvain resolutions (a) and numbers of neighbors (b) in the human liver data. a** The Louvain resolution varies from 0.3 to 0.7. The other parameters are $\alpha = 0$ and $r = 1.3$ which corresponds to that the mean number (around 4) of neighborhoods. **b** The radius varies from 1.3, 1.7 to 2.1 corresponding to the mean number of neighbors varying from 4 to 12. The other parameters are the Louvain resolution 0.6 and $\alpha = 0$.

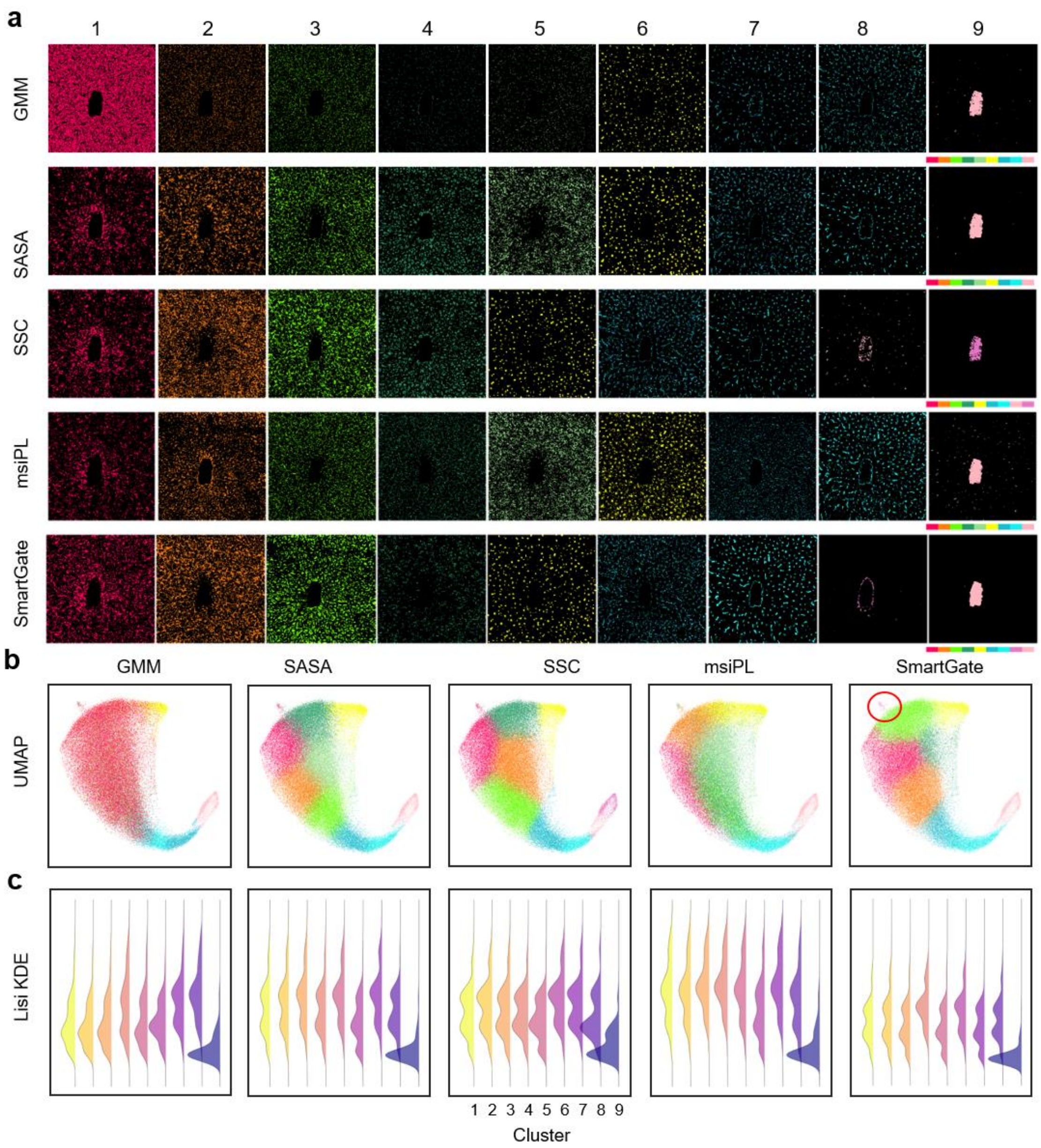


**Fig. S11. Comparisons of the spatial structures by SmartGate and other four methods in the mouse liver data. a** Each spatial structures identified by GMM, SASA, SSC, msiPL and SmartGate with the Louvain clustering. **b** The UMAP plots of spatial structures by the five methods respectively. **c** The kernel density plot of the LISI values for each structure of the five methods.

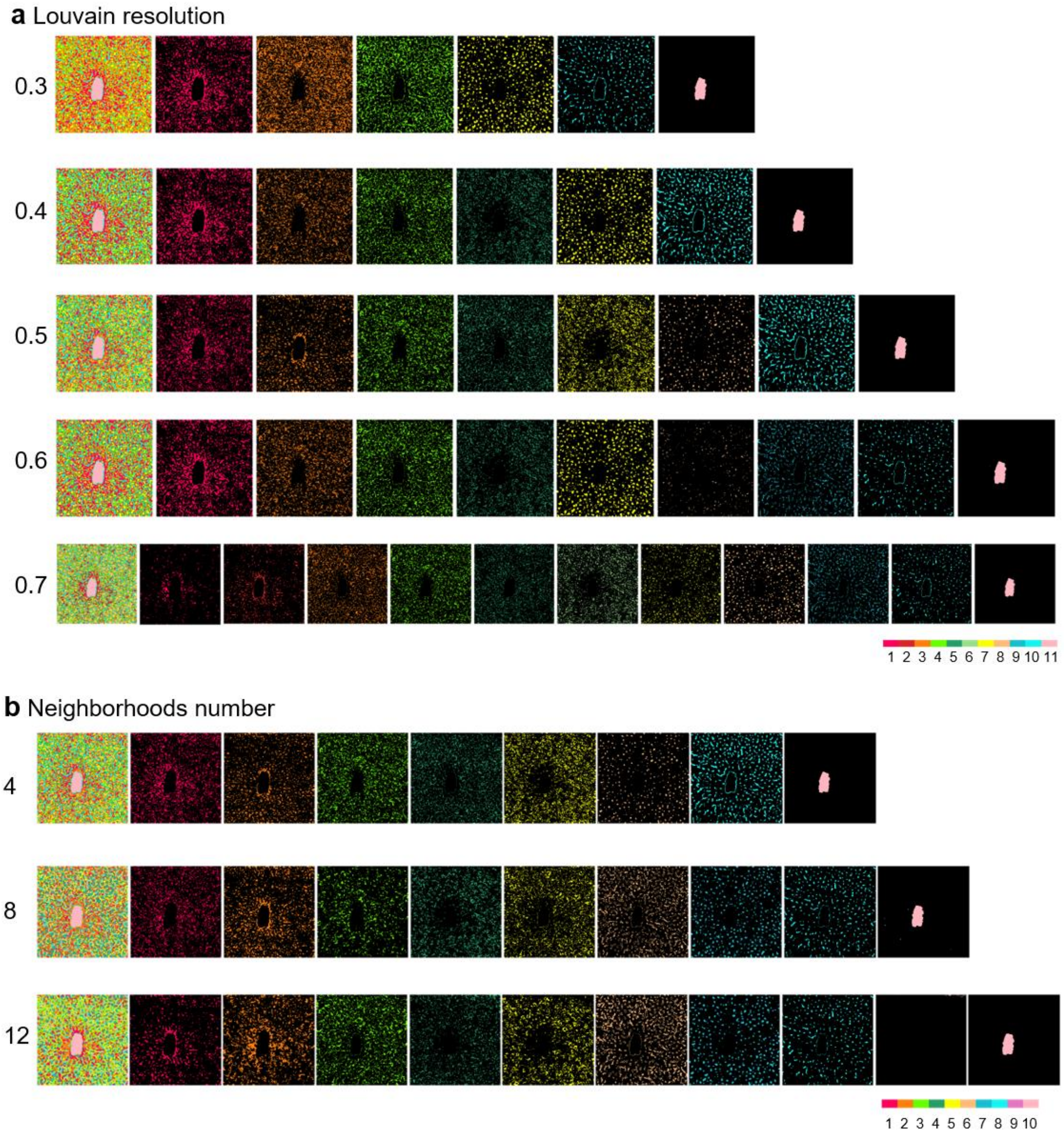


**Fig. S12. Comparison of spatial structures identified by SmartGate with diverse Louvain resolutions (a) and numbers of neighbors (b) in the mouse liver data. a** The Louvain resolution varies from 0.3 to 0.7. The other parameters are $\alpha = 0$ and $r = 1.3$ which corresponds to that the mean number (around 4) of neighborhoods. **b** The radius varies from 1.3, 1.7 to 2.1 corresponding to the mean number of neighbors varying from 4 to 12. The other parameters are the Louvain resolution 0.5 and $\alpha = 0$.

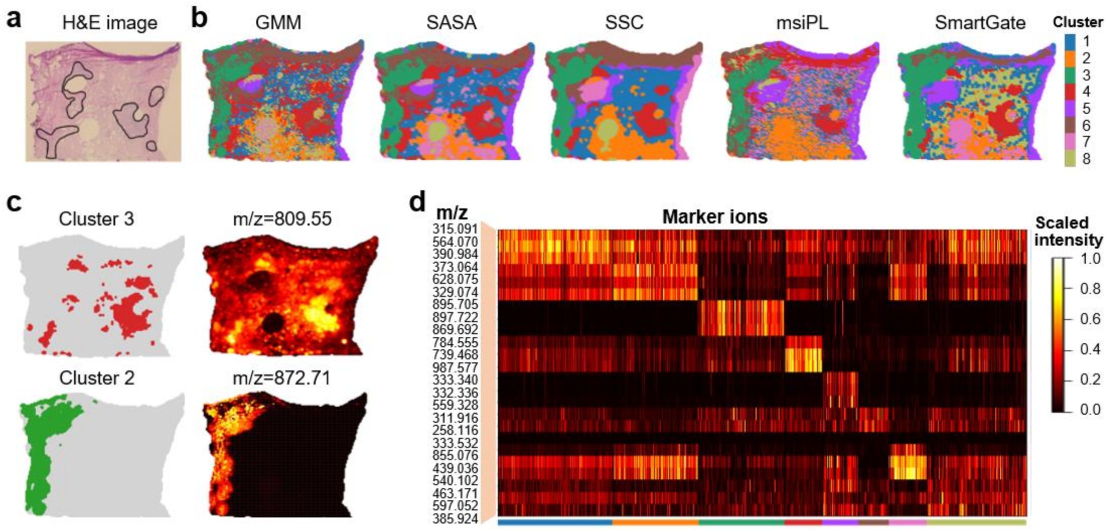


**Fig. S13**. **SmartGate identifies the main structures in the human prostate cancer dataset. a** The H&E image of the human prostate cancer. **b** Spatial structures identified by GMM, SASA, SSC, msiPL and SmartGate respectively. **c** Spatial distribution of two structures (clusters 2 and 3) and their corresponding marker ions (809.55 and 872.71) identified by SmartGate. **d** Top marker ions determined by differential analysis of each structure identified by SmartGate.

#### Supplementary Table

**Table S1. Main hyperparameter setting of SmartGate used in this study.**

| Dataset | Cell-type aware $\alpha$ | Pre-cluster Louvain resolution (if $\alpha \neq 0$) | Pre-defined radius $r$ | Embedding-cluster Louvain resolution or the number of components of mclust† | Whether automatic peak picking |
|---|---|---|---|---|---|
| Pig fetus | 0.5 | 0.1 | 1.4 | 10 | Yes |
| PDX mouse brain model of glioblastoma | 0.2 | 0.3 | 1.7 | 6 | Yes |
| 3D PDX mouse brain model of glioblastoma | 0.24 | 0.36 | 1.7 | 6 | Yes |
| Mouse kidney by nano-DESI | 0.3 | 0.2 | 1.3 | 0.3 | No* |
| Mouse kidney by MALDI | 0.3 | 0.2 | 1.8 | 9 | Yes |
| 3D Mouse kidney by MALDI | 0.3 | 0.2 | 2.6 | 9 | Yes |
| Human prostate cancer | 0.5 | 0.2 | 1.3 | 0.3 | No* |
| Human liver | 0 | 0.6 | 1.7 | 0.65 | No* |
| Mouse liver | 0.5 | 0.6 | 1.3 | 0.6 | No* |

†After automatic peak picking, we use mclust algorithm to cluster the embedding peak features. If not, we use Louvain algorithm to cluster.

*For these datasets, we only obtained the selected peaks (about 140~250 peaks) for further analysis without automatic peak picking.
